# Nanomechanics of Mandible Wear in Insects

**DOI:** 10.64898/2026.06.12.731969

**Authors:** Dilanka I. Deegala, Jonathan G. Pattrick, David Labonte

## Abstract

Many animals rely on specialised mouthparts to process food. Because this is a mechanically demanding task, mouthparts often wear, with potentially serious consequences for feeding performance and thus fitness. The biomechanics of wear are therefore of clear biological relevance, but remain poorly understood, especially in insects, where conventional engineering wear tests are hard to implement. Here, we present a nanomechanical characterisation of the mandibular epicuticle of three insect species: two leaf-cutting specialists, one with and one without transition-metal inclusions, and an omnivore. Contrary to predictions from simple engineering wear theory, wear resistance was neither directly proportional to indentation hardness nor inversely proportional to wear load. We suggest that this discrepancy arises in part from the high hardness-to-modulus ratio of mandibular epicuticle, which renders indentation hardness a poor proxy for resistance to plastic deformation. A simple elasto-plastic wear model qualitatively captures the main discrepancies between experiment and theory, and points to a revised set of wear proxies that may allow at least a qualitative ranking of biological materials via iso-performance lines on Ashby plots. Yet, as with most wear models, the wear coefficient remains unpredictable, a limitation strikingly illustrated by the increase in epicuticular wear resistance upon hydration despite a decrease in both hardness and modulus. Together, these observations suggest that purely plastic wear models may often be inadequate for biological materials with a high hardness-to-modulus ratio, and that even elasto-plastic models require careful validation against experimental wear assays.

**Graphical Abstract:** 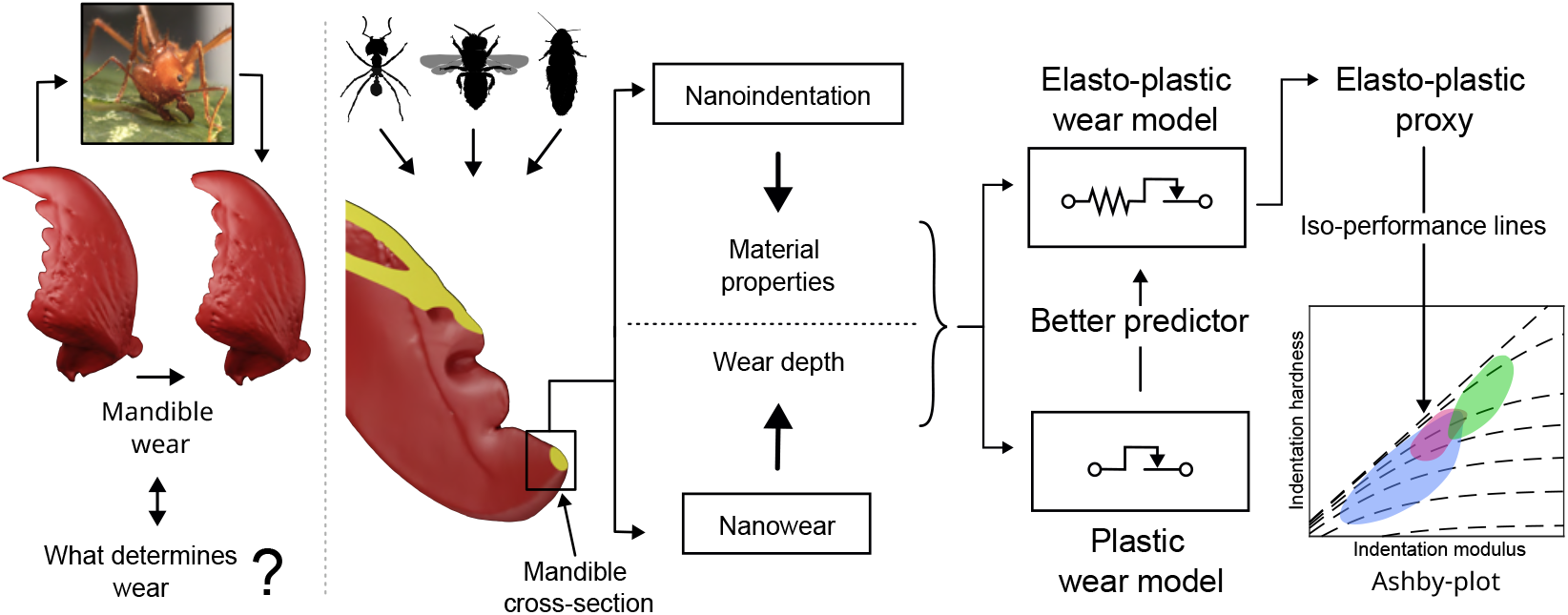

## 1. Introduction

Many animals use specialised mouthparts to mechanically break down food, thereby improving digestion and nutrient absorption efficiency [1, 2]. This is a mechanically demanding task that can involve high stresses and thus lead to removal of mouthpart material—mouthparts wear [3, 4]. To mitigate such wear and maintain functionality over many loading cycles, mouthparts often incorporate specialised hard tissues. Vertebrate teeth, for example, have a protective veneer of enamel, which is generally considered to be the hardest vertebrate tissue [5, 6]; and, likewise, mouthpart cuticle is typically the hardest and stiffest part of the arthropod exoskeleton, achieved through a locally increased chitin–protein crosslink density [7, 8] and, in some cases, the incorporation of reinforcing transition metals such as zinc or manganese [9, 10, 11, 12, 13, 7, 8]. Remarkable as these materials may be, they do not eliminate wear altogether, and, in fact, it often remains so pronounced that it has severe lifetime consequences in vertebrates [14, 15, 16, 17], and in invertebrates, where it reduces fecundity [18], larval growth [19, 20, 21], feeding rates [22, 18, 23, 24], pollination efficiency [25, 26], and life expectancy [27, 28]. Some have even gone so far as to suggest that repeated moulting in insects is an adaptation specifically to limit the adverse effects of wear [29, 30].

In light of the evidently severe biological consequences, mouthpart wear and the adaptations that may mitigate it have attracted considerable attention [6, 31, 32, 33, 34, 35, 36, 37, 20, 19]. Direct wear measurements nevertheless remain rare, especially for arthropod mouthparts [38, 12, 39, 40, 19, 27], and our understanding of the factors governing their wear therefore remains fragmentary [12, 41, 42, 43, 44, 45, 46, 47, 48, 49, 50, 51, 52, 53]. This is partly a technical problem: direct and controlled wear measurements are difficult to implement for small and often structurally heterogeneous arthropod mouth-parts [12, 39]. But it also reflects a theoretical challenge: wear is a complex system property that depends not only on material properties, but also on surface topography, contact conditions, environmental variables, and probabilistic effects that remain difficult to predict even in far more well-defined and homogeneous engineering materials [54, 55, 56].

A common response to these difficulties is to invoke simple wear proxies derived from contact-mechanics models of varying complexity, and expressed solely in terms of material properties that are easier to measure [57, 39, 9, 58, 59, 60, 61, 62, 63, 64, 65, 66]. Among the simplest, and to this day most widely used models is Archard’s law [67, 68, 69, 70], originally developed for sliding wear in metallic contacts [71]. The main idea is straightforward: the external work supplied during sliding is proportional to the product of normal load, *P*, and sliding distance, *s*; the work required to remove a unit volume of plastically deforming material in turn scales with material hardness, *H*. The wear volume, *V*, therefore is expressed as:

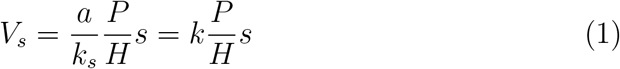

Where *k* is a dimensionless proportionality constant, often referred to as the wear coefficient. All else being equal, Archard’s law thus predicts lower wear volumes in harder materials, and this prediction has repeatedly underpinned functional interpretations of the mechanical performance of insect mouthparts in particular [9, 58, 59, 60, 61, 64, 65, 66]. More recently, indentation theory-derived property ratios, such as *H*^3^*/E*^2^ and *H/E*, have been proposed as alternative wear proxies across biological materials [57, 39, 63, 62]. What remains rare, however, is direct evidence that such proxies—whether based on hardness alone or on combinations of hardness and modulus—actually predict experimental rankings of wear resistance; and indeed, the limited evidence available suggests that they may not universally do so [72, 12].

Against this background, we here directly quantify the nanowear resistance of dry and hydrated mandibular epicuticle in three insect species, relate these measurements to nanoindentation-derived material properties, and test the extent to which wear models can account for experimentally observed wear patterns.

## 2. Materials and methods

### 2.1. Study animals

Mandibles from three species were used in both nanoindentation and nanowear experiments (Fig.1): *Atta cephalotes* (Linnaeus, 1758) leafcutter ants, chosen because they use their mandibles to mechanically process plant tissue at an almost industrial scale [73, 74, 75] and therefore are likely to experience strong selective pressure for increased mandibular wear resistance [11, 22, 47, 52]; leaf-cutter bees, *Megachile ligniseca* (Kirby, 1802), selected because, like leafcutter ants, they specialise in mechanical leaf-processing, but, unlike leafcutter ants, lack transition-metal incorporation at the mandible cutting edge [11, 12], offering an opportunity to test whether alternative strategies for increasing wear resistance exist; and Madagascan hissing cockroaches, *Gromphadorhina portentosa* (Schaum, 1853), included because the cockroach feeding apparatus has been identified as suitable model for that of generalist biting-chewing insects [76].

**Figure 1:**
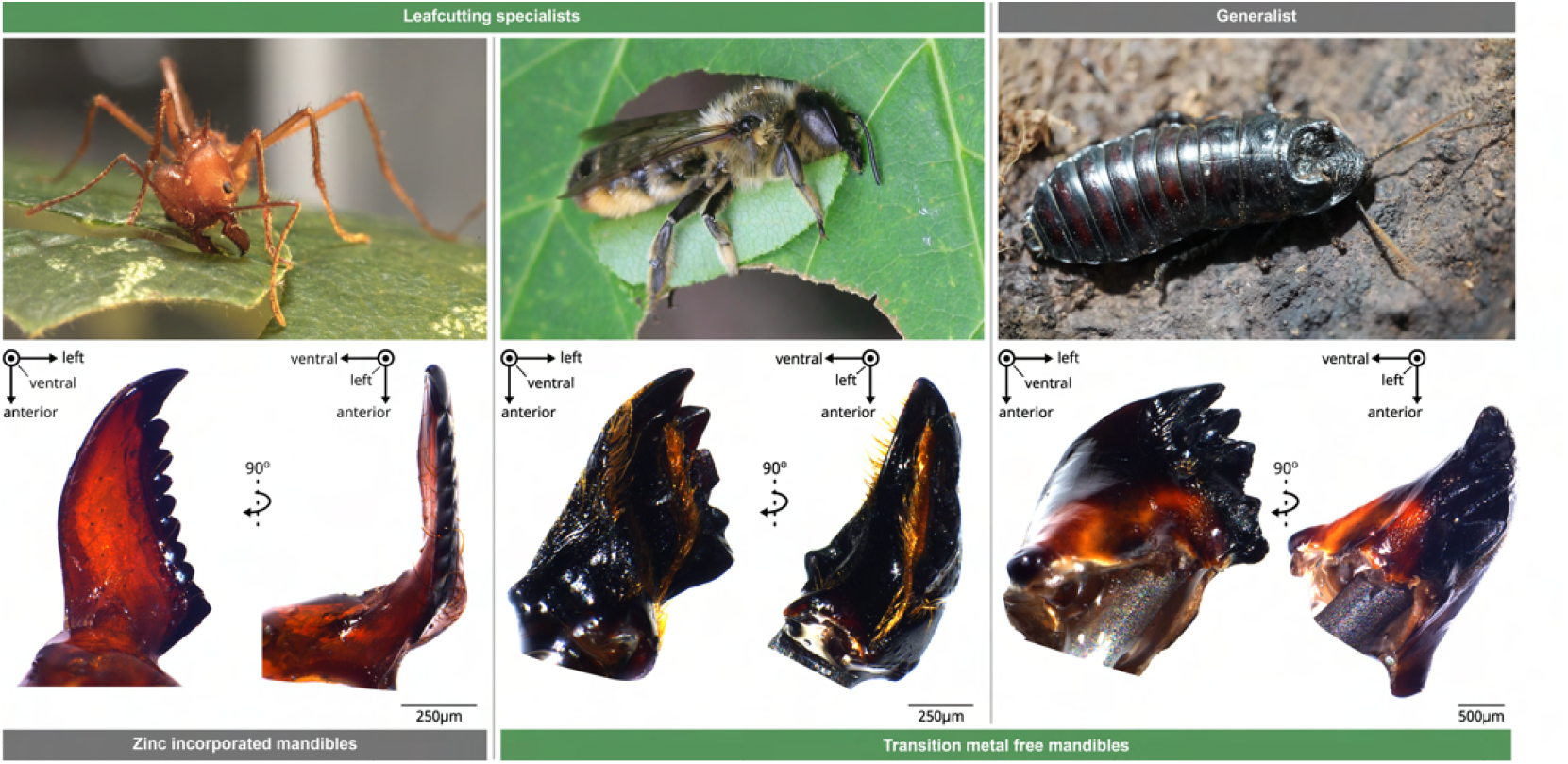
Three insect species used in this study. Top-left: Leafcutter ant, top-middle: leafcutter bees, top-right: hissing cockroach, bottom: ventral(left) and lateral(right) views of the mandibles.

Leafcutter ants and hissing cockroaches were collected from mature laboratory colonies kept in a climate chamber (FitoClima 12.000 PH, Aralab, Rio de Mouro, Portugal) at 25*°*C, 50–60% RH, and a 12/12 h light-dark cycle. Leaf-cutter ants were provided bramble, laurel, maize, and honey water; cat food, apples, and strawberries served as *ad libitum* food sources for hissing cockroaches. Leaf-cutter bees were collected in Oxfordshire and Worcester-shire between August and September in 2023.

Mature leaf-cutter ant workers vary by about two orders of magnitude in body mass. We used relatively large workers (50-60 mg body mass), because the thicker cuticle made testing easier. Leafcutter bees weighed 70-120 mg, and hissing cockroaches 6-8 g; in both cases, body size was dictated by availability. All insects were female.

### 2.2. Sample preparation

Insects were sacrificed by freezing, which has a negligible effect on the material properties of sclerotised cuticle [77, 78], and subsequently weighed to the nearest 0.1 mg (Explorer Analytical EX124, max. 120 g *×* 0.1 mg, OHAUS Corporation, Parsippany, NJ, USA). To facilitate mandible removal, thawed specimens were decapitated with a scalpel blade, and two ventral incisions were made to gain access to the mandibular apodemes, which hold the mandibles in place. With these apodemes cut, mandibles could then be prised loose with tweezers using little force. Any soft tissue remaining on the mandibles was carefully removed through gentle scrubbing with a scalpel blade.

Mandibles from nine individuals per species were selected for testing. Because leafcutter ant and bee mandibles are bilaterally symmetric [25, 26, 79], and preliminary nanomechanical tests failed to reveal evidence for noticeable differences in material properties between left and right mandibles (supple-mentary S1), one mandible per individual was randomly selected. Mandibles of cockroaches, in turn, are morphologically asymmetric [80, 76]; the right mandible was selected, because the larger number of teeth eased subsequent sample preparation (see below).

As is typical for fibre composites, arthropod cuticle can be anisotropic [81, 82, 83, 84], i. e., its mechanical properties may depend on the direction of measurement. To ensure consistent orientation among samples from the same species—and to realise a biologically meaningful and thus informative testing direction across species—custom jigs were designed (Fig.2). These jigs, manufactured via rapid prototyping (Form 3 resin printer, Formlabs), provided anchor points to which mandibles were glued: one anchor point for each of the two joint condyles, and two further points for the distal area of the external mandibular face and the basal margin, respectively [85, 86, see supplementary S2]. The orientation of these anchor points was species-specific and designed to control the orientation of the polished cross-section such that it was approximately normal to the bite force vector. The force applied during nanoindentation and wear assays is then approximately aligned with the bite force vector—a biologically relevant loading condition. Following previous work, we estimated the orientation of the bite force vector as the cross-product between the axis of mandible rotation and an arbitrary vector connecting this axis with the tip of the apical mandible tooth; the axis of rotation was defined as the vector passing through the centre of the two joint condyles [87, 88, 48]. To ease handling during subsequent sample preparation, the jigs were glued in pairs onto 13 mm circular glass coverslips (Stick2 Superglue HV, Everbuild building products LTD, UK)(See Fig.2 (b)).

**Figure 2:**
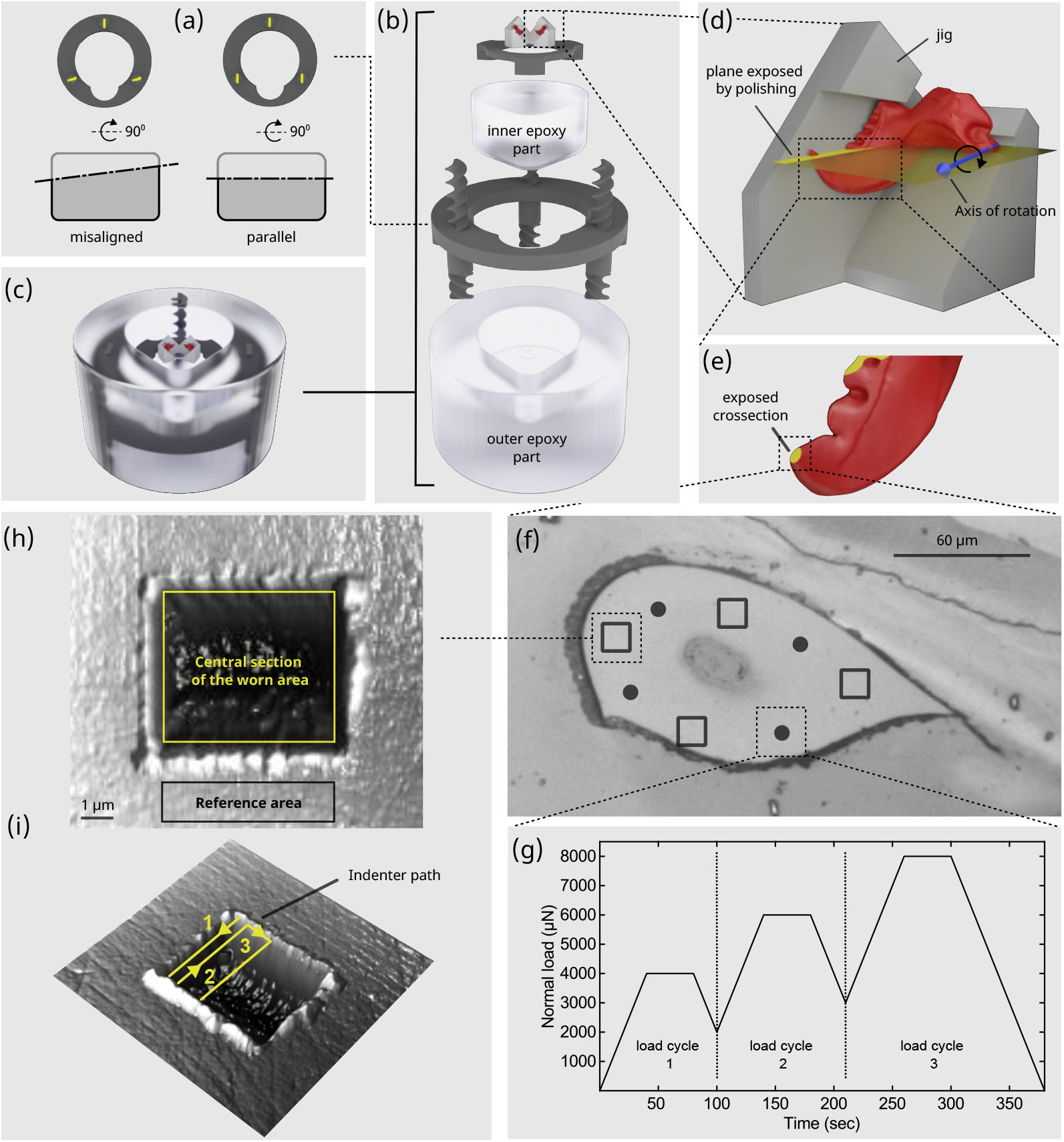
Sample preparation and mechanical characterisation. Insect mandibles were embedded in epoxy, followed by mechanical grinding and polishing to expose cross-sections of the apical tooth with defined orientation. Nanoindentation and wear tests were conducted on the epicuticle, readily identified by the brighter appearance compared to the darker inner layers. (a) The long axes of exposed spiral cross-sections are aligned only if the sample is polished evenly; sample pitch leads to spiral cross-section misalignment. The pitch of the spirals was chosen to achieve a plane misalignment of no more than 1*°* relative to the target plane. (b) Exploded view of the sample assembly. Top to bottom: mandibles - were glued to jigs and attached to cover slips in pairs; an inner component designed to fit the sample stage of the indenter stabilised the mandibles mechanically during polishing and testing; 3D-printed spirals helped ensure consistent sample orientation; and an outer component held the inner component in place during polishing. (c) Assembled sample. (d) To ensure a consistent and biologically relevant sample orientation, mandibles were attached to custom-designed jigs, fabricated by 3D printing. Each jig was designed such that the plane of the cross-section exposed during polishing was approximately in plane with the axis of rotation. (e) Schematic view of the exposed cross-section of the apical tooth of a leafcutter ant mandible, prepared for nanoindentation and wear experiments. (f) Microscopic image of the exposed epicuticle of a leafcutter ant mandible. Circles and squares indicate locations of nanoindentation and wear tests, respectively, separated by at least 10 µm in all tests. (g) A three-step load function was used to extract epicuticle indentation hardness and modulus as a function of depth. (h) After each wear treatment, a large surface scan at a low load revealed the surface topography of worn and unworn regions; the height difference between the unworn reference area and the central section of the worn region—the wear depth— served as an inverse proxy for the wear resistance. (i) Wear assays consisted of surface area scans at elevated normal loads. The arrows show the path followed by the indenter during the scan, which traced the surface in 256 lines at 3 Hz.

To mechanically stabilise the mandibles during grinding and polishing, the coverslips were next embedded within a two-component epoxy (epo-flow, Metprep, Coventry, UK), gently mixed and poured by hand, and cured at ambient conditions to minimise shrinkage. At this stage, one further constraint was imposed by the nanoindenter itself. A custom sample stage, used to control relative humidity during testing, fits up to four samples, each no larger than 5 mm *×* 15 mm (height *×* diameter). Each sample surface is acces-sible only through a circular hole, restricted to a 5 mm diameter to maintain stable experimental conditions inside the chamber via the Venturi effect. Because these size-constraints were incompatible with the standard fixtures of the machine used for grinding and polishing (Saphir 250 A2-ECO, ATM GmbH, Mammelzen, Germany), we designed two epoxy moulds: an internal component small enough to fit within the custom stage, but large enough to hold the jigs (15 mm diameter, with a small protrusion that helped orient the samples inside the environment control stage); and an external component with a diameter of 30 mm, and a small internal cavity that holds the internal component in place during polishing (see Fig.2(b-c)). A custom-designed and resin-printed part, consisting of three spirals with a rectangular crosssection, was embedded in the external component, to help maintain sample orientation during polishing: only if the samples were polished evenly would the long axes of all three spirals appear aligned in the exposed cross-section (Fig.2(d)). The spirals were designed with a pitch of 3.6 mm; a long axis deviation angle of approximately 10*°*, easily identifiable by the naked eye, then corresponds to a 0.5*°* tilt in the sample surface, defining the approximate accuracy of this method.

Mandible mechanical properties may vary not only with testing orientation, but also along the long axes of the exposed teeth (Fig.2(e-f)). To obtain a rough estimate for the magnitude of such variation, we conducted a preliminary study in which leafcutter ant mandibles were repeatedly polished and tested; although the effect size was small, material properties appeared to systematically decreased with depth (supplementary S3). To reduce the variation stemming from such variation, all samples were polished such that the exposed cross-section of the distal-most tooth had a long-axis width of about 100 µm. For polishing, samples were first mechanically ground using abrasive paper of increasing fineness (P800-P4000, Metprep, Coventry, UK), and subsequently polished using 0.3 µm polishing alumina dispersed on polishing cloths. This sequence resulted in a consistent average roughness, *R*_*a*_, of no more than 40 nm (measured via scanning probe microscopy (see next section))

To standardise sample hydration,polished samples were stored in sealed containers filled with drying pearls for at least 6 hours prior to mechanical characterisation (Drying pearls orange, Sigma-Aldrich).

### 2.3. Mechanical characterisation

With cross-sections of controlled orientation exposed, the next task was to design robust protocols for the characterisation of both cuticle material properties and wear resistance—a system property. Both types of tests shared three methodological elements.

First, all experiments were conducted on a Hysitron TI 980 Triboindentor (Bruker, MN, USA). Material property measurements were conducted with a diamond Berkovich xSol25 tips (TI-0283, Bruker) and a standard 1D transducer, whereas wear assays used conospherical diamond xSol25 tips (TI-0040, Bruker) with 300 nm radius and a standard 2D transducer; they thus loosely mimicked mandible wear induced by small hard particles such as phytoliths [20, 27]. Both transducers were connected to the granite base of the Triboindenter via a piezo scanner (TriboScanner, Bruker). Tip area functions were determined according to manufacturer instructions, and took the form 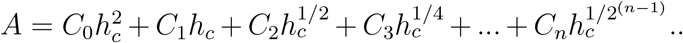 where *C* are constants, and *h*_*c*_ is the contact depth. For Berkovich tips, *C*_0_ was fixed at the theoretical expectation of 24.5, and *C*_1_ *− C*_3_ were fitted as higher-order correction terms; for conospherical tips, *C*_0_ was fixed at *π*, and *C*_1_ *− C*_4_ were fitted (supplementary S4). Tip area calibration was performed once before experiments for each of the three species, and remained highly consistent throughout, indicating minimal tip wear; system compliance was calibrated once a year.

Second, all mechanical testing was performed on the distal-most tooth, and specifically its epicuticle, because it represents the first barrier against wear, and because its amorphous structure reduces the likelihood of anisotropy. The epicuticle can be readily identified by its smooth finish, which reflects more light and appears brighter than the layered sections of the exocuticle (Fig.S6-S9); indentations and scratch tests confirmed this visual classification by revealing consistent differences in mechanical response (see [89] and supplementary S5).

Third, recognising that the mechanical properties of insect cuticle can depend strongly on hydration [39, 11, 7, 90, 63, 84, 38, 91, 92, 93, 94, 95], both nanoindentation and nanowear measurements were conducted at two relative humidities (RH) selected to represent dry (30%) or hydrated conditions (90%), respectively. Preliminary experiments suggested that material property variation was negligible once RH dropped below 90%, and 30% was a conservative choice at the lower limit of the environmental chamber (supplementary S6). Local RH was maintained within the small volume of a custom-made sample stage—modified by Bruker from their standard xSol environmental control stage— via a humidity generator (HGC 30, Data-Physics Instruments), which supplied humid air at a maximum flow rate of 200 mL s^*−*1^. Prior to all experiments, samples were placed inside the chamber for at least 6 hours to equilibrate, a time frame informed by preliminary tests following Klocke et al. [84, See supplementary S7].

At 90% RH, indentation hardness and modulus did not decrease significantly over a period of 4 hours following the 6-hour equilibration period, further indicating that 6 hours was sufficient for the epicuticle hydration to reach a steady state (See supplementary S8).

#### 2.3.1. Nanoindentation

Three epicuticle material properties were measured via nanoindentation in load control: indentation modulus, indentation hardness, and “true” hardness (see data analysis and statistics). As indentation sites were identified from optical images of the exposed cross-sections, the thickness of the exposed epicuticle could not be determined directly; where it is comparable to the indentation depth, mechanical properties measurements may become unreliable, identifiable by variation of mechanical properties with indentation depth [96, 84]. To probe for such variation, we used a partial unloading load function (also see [97, 98, 99]) with three load cycles, representing a compromise between the need for short measurements to manage drift on one hand and depth resolution on the other (Fig.2(g)). The lowest load was chosen to ensure that indentation depths exceeded 10*R*_*a*_ *≈* 400 nm, required to reduce the effects of surface roughness [100, 101]; the highest load was close to the maximum load capacity of the transducer; and the intermediate load was half-way in-between, leading to loads of 4000, 6000, and 8000 µN for leafcutter ant and bee mandibles, and of 2000, 4000, and 6000 µN for hissing cockroach mandibles, which preliminary trials revealed as more compliant. Any indentation trials that revealed a discontinuity in property variation with depth were qualitatively identified and excluded; properties were compared at maximum load to maximise the signal-to-noise ratio.

Each load cycle consisted of three steps. The load was ramped up at a low rate of 100 µN s^*−*1^ to minimise viscoelastic effects [84, 39, 102], typically small for sclerotised cuticle to begin with [103]. To eliminate nosing, it was then kept at the maximum until the creep rate had dropped to levels comparable to the transducer drift rate (about 1 nm s^*−*1^). For mandibles of leafcutter ants and bees, this requirement was met by hold times of 40 and 65 s at 30% and 90% RH, respectively, while for hissing cockroach mandibles, 65 s were required. Partial unloading was then conducted at 100 µN s^*−*1^, limited to 50% of the preceding maximum load to ensure that the indenter tip remained in surface contact throughout [104, 105].

Using this loading protocol, we measured the mechanical properties of the epicuticle at four locations per sample for each hydration, resulting in a total of 216 indents. Indents were placed randomly but such that they were separated from other test locations and the sample’s edge by at least 10 µm—about ten times the contact depth, as required to avoid interaction artefacts [105, 106].

#### 2.3.2. Nanowear assay

A dominant source of insect mandible wear is thought to arise from abrasion by hard particles such as phytoliths [37, 107, 19, 108, 27, 109], and to loosely simulate this process, we conducted nanowear tests. During each wear test, the indenter tip is used in scanning probe microscopy mode to raster across the sample surface, resulting in localised removal of material [39, 102, 89]. To quantify the resulting wear, surface topography scans are then conducted at a lower load after each wear treatment.

Nanowear tests were conducted in load control mode at normal loads of 20, 40, 70, and 100 µN. Lower loads were not used as that would require a large number of passes (see below), drastically increasing the test time. Loads higher than 100 µN, in turn, result in depths exceeding the approximately 100 nm sphere-to-cone transition depth of the indenter (supplementary S9). Each individual wear treatment consisted of multiple passes across an area of 6 µm *×* 6 µm, and each pass consisted of 256 line scans (see Fig.2(h)), conducted at 3 Hz. Preliminary trials confirmed that the relationship between wear depth and the number of passes was approximately linear (supplementary S10). The benefit of this observation was that it permitted increasing the number of passes at low loads to ensure that wear exceeded the noise floor defined by the surface roughness, and reducing number of passes at high loads, to avoid excessive wear; 25, 21, 5 and 3 passes were used for 20, 40, 70, and 100 µN wear trials, respectively, determined by trial and error. Surface topography scans were conducted at 2 µN after each wear treatment. A larger scan area of 12 µm *×* 12 µm ensured that both the worn and unworn reference regions were included, permitting quantification of the material removed—an inverse proxy for material wear resistance [39, 110, see data analysis and statistics].

Four randomly placed wear tests, each with a different normal load, were conducted per sample at each hydration level, leading to 216 tests across all samples. A minimum distance of 10 µm was maintained between the boundaries of wear tests, indent locations and the sample edge to minimise interaction artefacts.

### 2.4. Visual and elemental analysis

We inspected the wear area with SPM (Scanning Probe Microscopy), optical microscopy and SEM (Scanning Electron Microscopy) to identify the wear patterns emerging from the wear tests. For a small number of additional test samples, post-wear surface scans were not conducted as they remove wear debris (Fig.6). Worn samples were first coated with a 20 nm Chromium layer (Quorum Q150TS), and then imaged using secondary electrons with an accelerating voltage of 3 kV.

Transition metals are present in the cuticle of many species of arthropods [39, 58, 111, 12], including leafcutter ants [11, 112]. To confirm the presence of transition metals expected from the literature [11, 112, 12], the element composition of randomly selected points on the epicuticle was measured via Energy Dispersive X-ray Spectroscopy (EDS) at an acceleration voltage of 10 kV: 16 points per mandible of leafcutter ants and bees, 10 points for hissing cockroach, and two mandibles per species. As expected, Zinc was the only transition metal detected by EDS; it was present in every location tested on the epicuticle of ant mandibles [12], but absent from bees [12] and cockroach mandibles.

### 2.5. Data analysis and statistics

Three mechanical material parameters were extracted from nanoindentation experiments: indentation hardness, *H*_*I*_, indentation modulus, *E*_*I*_, and true hardness, *H*. Indentation hardness is defined as the mean contact pressure at maximum load; it is therefore an elasto-plastic hybrid parameter that reflects the resistance of the material to both plastic (permanent) and elastic (reversible) deformation [113, 114, 115, 116]. The resistance to elastic deformation, in turn, is quantified by the indentation (or reduced) modulus. In indentation, it is defined strictly as the inverse of the sum of the plane-strain elastic compliances of the indenter and the sample, respectively [114]. Because diamond, the indenter material, has a much higher plane-strain modulus than the epicuticle, the indentation modulus is practically equal to the epicuticle plane-strain modulus. The resistance to irreversible deformation, then, is quantified by the true hardness, *H*, represented conceptually by a plastic slider [116, 117, 62]; *H* may be interpreted as the mean contact pressure during the indentation of a rigid-perfectly-plastic material, or, equivalently, as the energy required to create a unit volume of purely irreversible deformation [116].

*H*_*I*_ and *E*_*I*_ were obtained from the force–displacement data using the Oliver–Pharr method [118]; *H* was then estimated using a lumped-parameter model of an ideal elasto-plastic material, formed by an elastic spring and plastic slider in series [116, 119, 62]. Indentation hardness was calculated as:

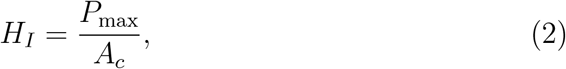

where *A*_*c*_ = *A*(*h*_*c*_) is the projected contact area between the tip and material at peak load. To extract this area from the tip area function, *h*_*c*_ was determined from the maximum normal load *P*_max_, maximum depth *h*_max_, and unloading stiffness *S*:

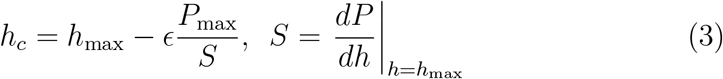

Here, *ϵ* is a geometric constant, equal to 0.75 for a Berkovich tip [114]. To obtain *S*, the tangent to the force–displacement curve at maximum load, a power-law function of the form *P* = *α*(*h−h*_*f*_)^*m*^ was fitted to the unloading segment between 95% and 20% of the maximum load, treating *h*_*f*_, *α*, and *m* as variables. Then the indentation modulus follows as:

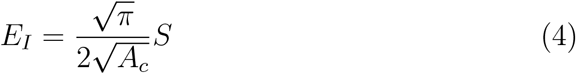

The true hardness is a sole function of *H*_*I*_, *E*_*I*_, and the indenter geometry, represented by *β*, the equivalent cone angle measured relative to the sample surface:

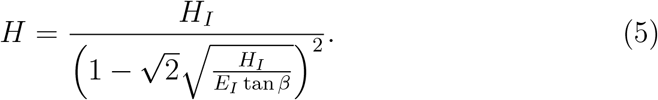

For a Berkovich tip, 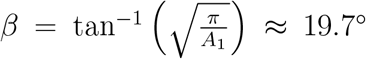. To investigate how well the ideal elasto-plastic material model approximates the mechanical behaviour of the mandible epicuticle, we used it to predict the residual-to-maximum indentation depth, *h*_*r*_*/h*_*max*_, to then compare it with the experimentally determined value, using t-test with Satterthwaite’s approximation (see section 3.1 and supplementary S13). For an ideal elasto-plastic material:

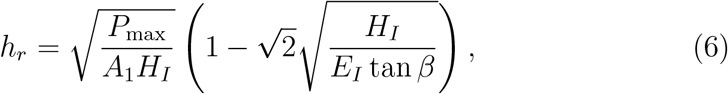

Wear area visualisation and wear volume quantification were completed in Gwyddion 2.62 [120]. Following tilt correction, a smooth and unworn reference area of at least 6 µm *×* 3 µm was selected, and the difference in mean height between the reference area and the central section of the worn area was defined as the wear depth *d*.

Two further corrections were necessary to account for systematic differences introduced by the variation in wear loads. First, *d* was divided by the total number of passes to retrieve a wear depth per pass. Second, even within one pass, a fraction of the material may undergo more than two effective wear cycles, because the distance between two scan lines is not necessarily equal to the contact diameter between the indenter tip and epicuticle. To correct for this effect, the number of effective wear cycles was estimated as the ratio between the contact diameter and the centre-to-centre distance between two scan lines. Combining both corrections leads to the definition of the normalised wear depth *W*_*h*_ per effective wear cycle:

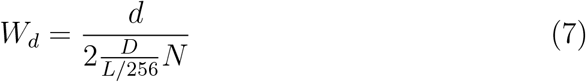

where *d* is the wear depth, *L* is the side length of the worn area,*N* is the number of passes, *D* is the contact diameter, 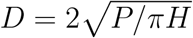, and the factor two arises because each scan line comprises a forward and backward trace.

To compare epicuticle indentation hardness, modulus, and true hardness across species, and to assess the effect of hydration on these material properties, we used two-way analysis of variance (ANOVA) with sample ID as the repeated measure; a three-way ANOVA with sample ID as the repeated measure assessed the effect of normal load, species, and RH on normalised wear depth. All ANOVAs reported use Type III sum of squares.

To test the validity of Archard’s law for insect epicuticle, Eq.(1) was rewritten to predict the normalised wear depth *W*_*d*_ (see supplementary S11).

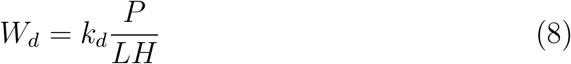

Here, *L* is the side length of the wear patch, and *k*_*d*_ is a dimensionless constant, related to Archard’s wear coefficient. Two predictions were then tested, using ordinary least squares (OLS) regression on log_10_-transformed data. First, we tested whether *W*_*h*_ and *H*_*I*_ are inversely proportional at each load level (slope of negative unity). Second, we tested whether *W*_*h*_ and *P* were directly proportional within species (slope of unity), where indentation hardness varied little (see results). Because statistical analysis revealed a significant interaction between wear load and RH (see Results), both tests were conducted separately for each RH-level. All statistical analyses were conducted in MATLAB R2024a (MathWorks, Natick, MA, USA), and values in the text indicate arithmetic mean *±* standard error unless otherwise indicated.

## 3. Results

### 3.1. Mandible material properties differ across species, and decrease with relative humidity

True hardness, indentation hardness and indentation modulus differed significantly across species (RM ANOVA Species, *H*: *F*_2,24_=91.47, *p<*0.0001, *H*_*I*_: *F*_2,24_=76.09, *p<*0.0001, *E*_*I*_: *F*_2,24_=14.59, *p<*0.0001), independent of RH-level (RM ANOVA Species *×* RH, *H*: *F*_2,24_=0.03, *p<*0.968, *H*_*I*_: *F*_2,24_=0.14, *p*=0.873, *E*_*I*_: *F*_2,24_=1.28, *p*=0.297. See Fig.3(a-b), Table 1, and supplementary S12). At both RH levels, *H, H*_*I*_ and *E*_*I*_ of leafcutter ant mandible epicuticle were about 45%, 25% and 9% higher than that of leafcutter bee and hissing cockroach mandibles respectively (Tukey-Kramer on average across RH, ants vs bees: *H*: *q*_24_=16.70, *p<*0.0001, *H*_*I*_: *q*_24_=15.12, *p<*0.0001, *E*_*I*_: *q*_24_=6.11, *p*=0.0007, ants vs cockroach: *H*: *q*_24_=16.43, *p<*0.0001, *H*_*I*_: *q*_24_=15.09, *p<*0.0001, *E*_*I*_: *q*_24_=7.03, *p*=0.0001), which did not differ significantly in their material properties at either RH (Tukey-Kramer on average across RH, bees vs cockroaches: *H*: *q*_24_=0.27, *p*=0.98, *H*_*I*_: *q*_24_=0.03, *p>*0.99, *E*_*I*_: *q*_24_=0.92, *p*=0.79). Increasing RH from 30 to 90% consistently decreased *H, H*_*I*_ and *E*_*I*_ by about 10% in all species (RM ANOVA RH, *H*: *F*_1,24_=23.01, *p<*0.0001, *H*_*I*_: *F*_1,24_=49.40, *p<*0.0001, *E*_*I*_: *F*_1,24_=74.56, *p<*0.0001).

**Figure 3:**
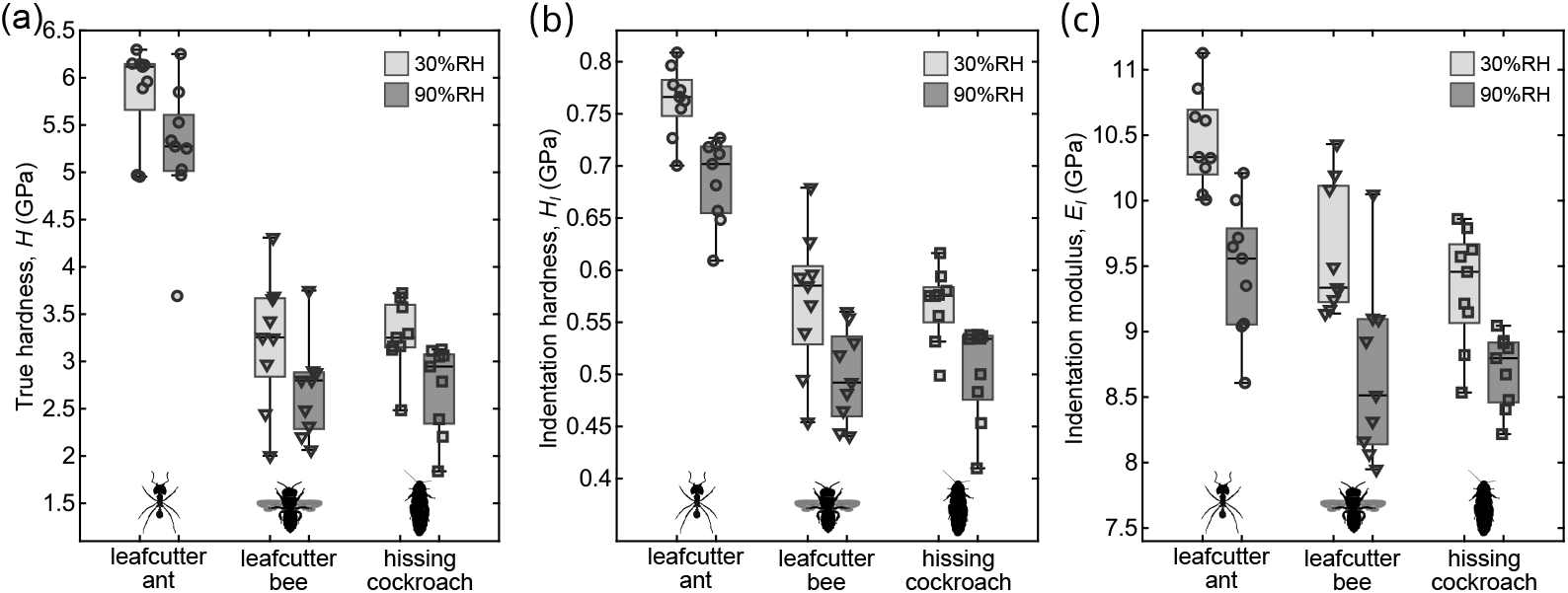
Results of nanoindentation. (a)-(c) True hardness, indentation hardness and indentation modulus were 45%, 25% and 9% higher in leafcutter ant mandibles, but did not significantly differ between leafcutter bee and hissing cockroach mandibles. An increase in relative humidity from 30-90% reduced both material properties by about 10% in all the mandibles. In all box plots, the central line indicates the median, the box bounds represent the 25th and the 75th percentile, and whiskers extend to the most extreme data points above and below the median that are within 1.5 times the interquartile range from the box bounds.

**Table 1:** True hardness, *H*, indentation hardness, *H*_*I*_, and indentation modulus, *E*_*I*_, of the epicuticle of leafcutter ant, leafcutter bee and hissing cockroache mandibles at 30% and 90% relative humidity (RH)

| species | RH | $H$ (GPa) | $H_I$ (GPa) | $E_I$ (GPa) |
| --- | --- | --- | --- | --- |
| Ant | 30% | $5.85 \pm 0.17$ | $0.76 \pm 0.01$ | $10.47 \pm 0.12$ |
| | 90% | $5.24 \pm 0.24$ | $0.69 \pm 0.01$ | $9.47 \pm 0.17$ |
| Bee | 30% | $3.22 \pm 0.23$ | $0.57 \pm 0.02$ | $9.60 \pm 0.17$ |
| | 90% | $2.69 \pm 0.17$ | $0.50 \pm 0.01$ | $8.69 \pm 0.22$ |
| cockr-<br>oach | 30% | $3.27 \pm 0.13$ | $0.57 \pm 0.01$ | $9.34 \pm 0.15$ |
| | 90% | $2.72 \pm 0.16$ | $0.50 \pm 0.02$ | $8.70 \pm 0.09$ |

The residual-to-maximum depth ratio predicted by the elasto-plastic model did not differ significantly from the values obtained from nanoindentation force displacement data (*t*_91.68_=1.23, *p*=0.22. See supplementary S13).

### 3.2. Wear depth differs across species, and increases with wear load and relative humidity

Increasing wear load was associated with an increase in normalised wear depth, *W*_*d*_ (RM ANOVA Load, *F*_3,72_=600.6, *p<*0.0001, Fig.4). This effect was strongest in leafcutter ant mandibles (RM ANOVA Species, *F*_2,24_=98.9, *p<*0.0001) i.e. there was a significant species-load interaction (RM ANOVA Species *×* Load, *F*_6,72_=21.3, *p<*0.0001): at a wear load of 20 µN, the normalised wear depth of leafcutter ant mandibles was a factor of about 3-4 lower than that of cockroach and bee mandibles; at a wear load of 100 µN, this difference decreased to a factor of 1.5-2. Notably, normalised wear depths of hissing cockroaches were about 1.2-2 higher than that of leafcutter bees at all loads (see Fig.4 and supplementary S12 for detailed statistics).

**Figure 4:**
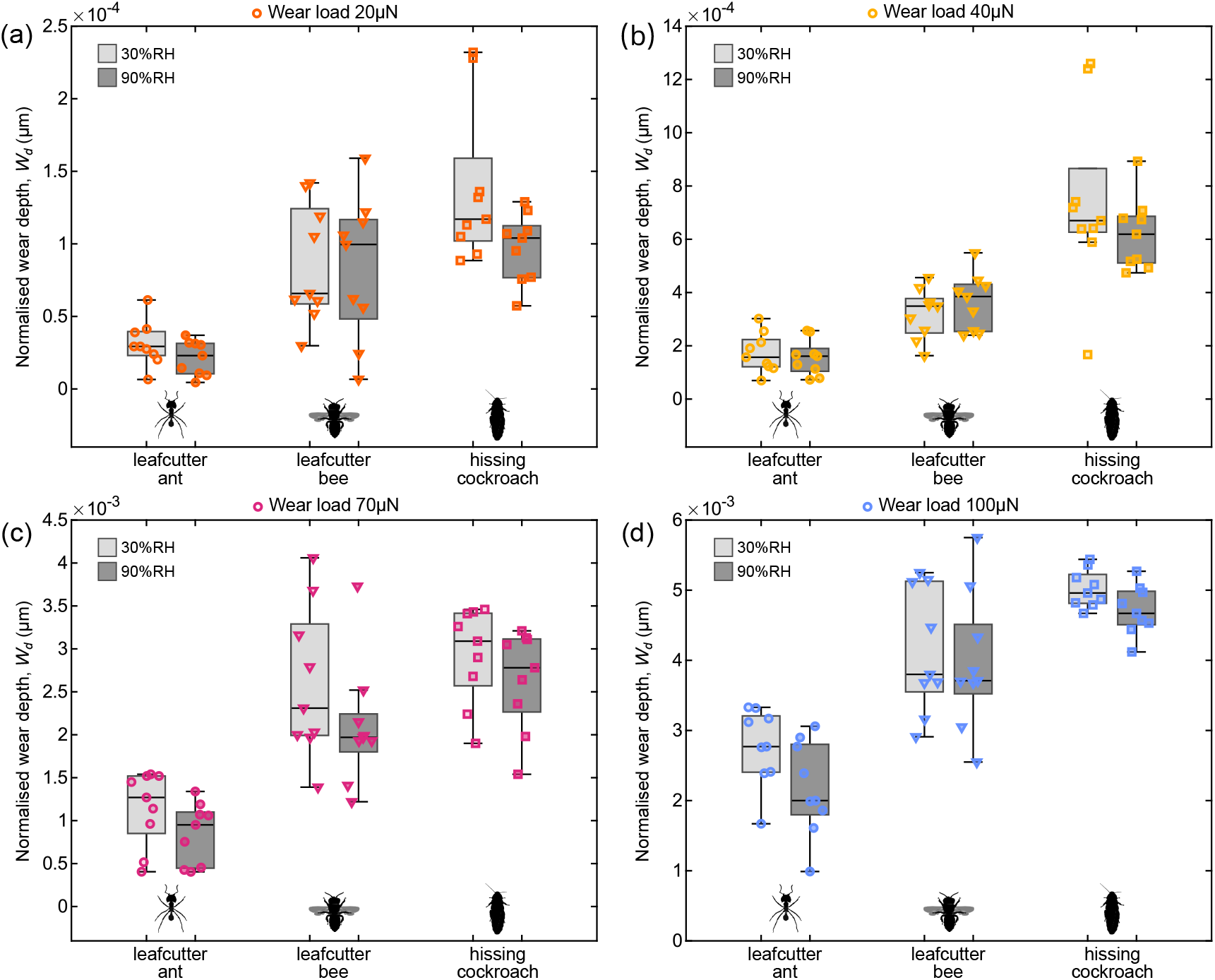
Results of nanowear tests. (a–d) Among the three species, normalised wear depth was lowest in leafcutter ant mandibles, but the extent of this difference decreased with wear load. At 20 µN, it was approximately 3–4 times lower than in hissing cockroach and leafcutter bee mandibles, whereas at 100 µN it was only 1.5–2 times lower. Increasing relative humidity from 30% to 90% reduced normalised wear depth by about 10%, although this effect was only significant at the two highest wear loads, 70 and 100 µN. In all box plots, the central line indicates the median, the box bounds represent the 25th and the 75th percentile, and whiskers extend to the most extreme data points above and below the median that are within 1.5 times the interquartile range from the box bounds.

An increase in RH was associated with a decrease in normalised wear depth by about 10% for all species (RM ANOVA RH, *F*_1,24_=9.11, *p*=0.0059). However, this effect was restricted to the two highest loads only (RM ANOVA RH *×* Load, *F*_3,72_=3.6, *p*=0.018 .Fig.3(c-f) and supplementary S12 for detailed statistics).

### 3.3. Variation of normalised wear depth with wear load and indentation hardness contradicts Archard’s law

In notable contradiction to Archard’s law, normalised wear depth did not generally vary in inverse proportion to hardness at either RH. Instead, power-law exponents were up to three times smaller at the lowest wear loads, *h ∝ H*^*−*3^, indicating a much stronger dependence of normalised wear depth on indentation hardness (Fig.5 (a-b) see Table 2). However, the gradient gradually approached the prediction of negative unity as wear loads increased, and was no longer statistically different from it at 100 µN (Fig.5 (a-b)).

**Figure 5:**
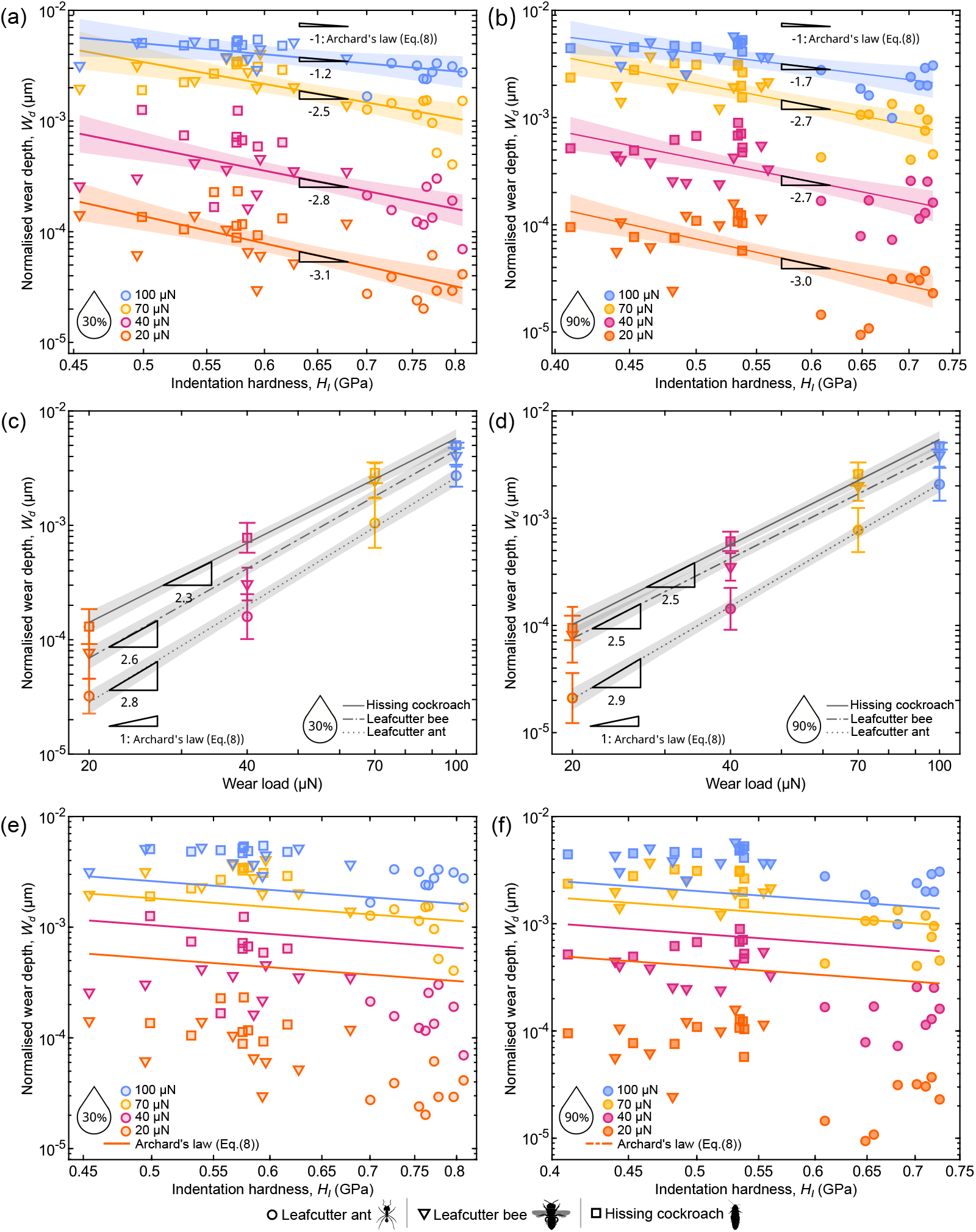
Evaluation of Archard’s law prediction against experimental data and Archard’s law fit to the experimental data. To test the simple predictions from Archard’s law, we conducted linear regression on log_10_-transformed normalised wear depth and indentation hardness at 30% (a) and 90% relative humidity (RH) (b) and extracted the power law - exponent. For both RH levels, this exponent is significantly smaller than Archard’s law prediction of negative unity for all lower loads except 100 µN. The power law exponent obtained from linear regression between log-transformed normalised wear depth and normal load at 30%RH (c) and 90%RH (d) is significantly larger than unity, the prediction from Archard’s law. The exponent was highest for ant mandibles with the highest indentation and true hardness. Archard’s law Eq.(8) was fitted to the experimental data obtained at all loads by estimating *k*_*d*_ at 30%RH (e) and 90%RH (f). See supplementary S12 for the fitted values. Error bars in (c) and (d) represent the standard deviation

**Table 2:** Results of mixed linear regression on log_10_-transformed normalised wear depth and indentation hardness at 30% and 90% relative humidity (RH). 95% confidence intervals are provided in parentheses.

| load(μN) | RH | gradient | $F_{1,197}$ | p-value |
| --- | --- | --- | --- | --- |
| 20 | 30% | -3.10 [-4.23, -1.98] | 13.58 | 0.0003 |
|  | 90% | -2.99 [-4.08, -1.91] | 13.14 | 0.0004 |
| 40 | 30% | -2.76 [-3.85, -1.67] | 10.12 | 0.0017 |
|  | 90% | -2.72 [-3.75, -1.68] | 10.75 | 0.0012 |
| 70 | 30% | -2.49 [-3.58, -1.40] | 7.30 | 0.0075 |
|  | 90% | -2.68 [-3.71, -1.64] | 10.23 | 0.0016 |
| 100 | 30% | -1.23 [-2.32, -0.14] | 0.18 | 0.6717 |
|  | 90% | -1.69 [-2.72, -0.66] | 1.73 | 0.1895 |

The relationship between normalised wear depth and load, too, was significantly steeper than a direct proportionality across all species, and instead followed a quadratic or even cubic dependency for either RH (Fig.5 (c-d) see Table 3). The deviation from a direct proportionality seemed strongest in leafcutter ant mandibles, which had the highest indentation hardness.

**Table 3:** Results of mixed linear regression on log_10_-transformed normalised wear depth and wear load for 30% and 90% relative humidity (RH). 95% confidence intervals are provided in parentheses.

| species | RH | $H_I$(GPa) | gradient | $F_{1,200}$ | p-value |
| --- | --- | --- | --- | --- | --- |
| ant | 30% | 0.76 | 2.82 [2.61, 3.02] | 307.56 | <0.0001 |
|  | 90% | 0.69 | 2.87 [2.66, 3.07] | 325.65 | <0.0001 |
| bee | 30% | 0.57 | 2.60 [2.40, 2.80] | 254.68 | <0.0001 |
|  | 90% | 0.50 | 2.48 [2.27, 2.68] | 204.06 | <0.0001 |
| cockr- | 30% | 0.57 | 2.30 [2.11, 2.50] | 168.56 | <0.0001 |
| oach | 90% | 0.50 | 2.47 [2.27, 2.67] | 214.54 | <0.0001 |

### 3.4. Dominant wear mode appears to be independent of wear load

Two distinct features of the wear areas were observed in SEM images, independent of the wear load, species and hydration (see Fig.6a-d). First, the worn surfaces remained relatively smooth, closely resembling unworn regions. Second, the wear debris consisted mainly of smooth-edged flakes, often agglomerated and associated with peripheral pile-up. The absence of surface cracks, scratch marks, ribbon-like debris, deep grooves, radial cracking, and angular chips indicates that the dominant wear process is unlikely to involve brittle fracture, abrasion, or abrasive micro-cutting [121, 122, 123]. Instead, the observations are most consistent with adhesive and delamination wear, in which repeated sliding contact promotes ductile/plastic deformation and smearing of the cuticle before local detachment of debris [124, 125, 126, 127].

**Figure 6:**
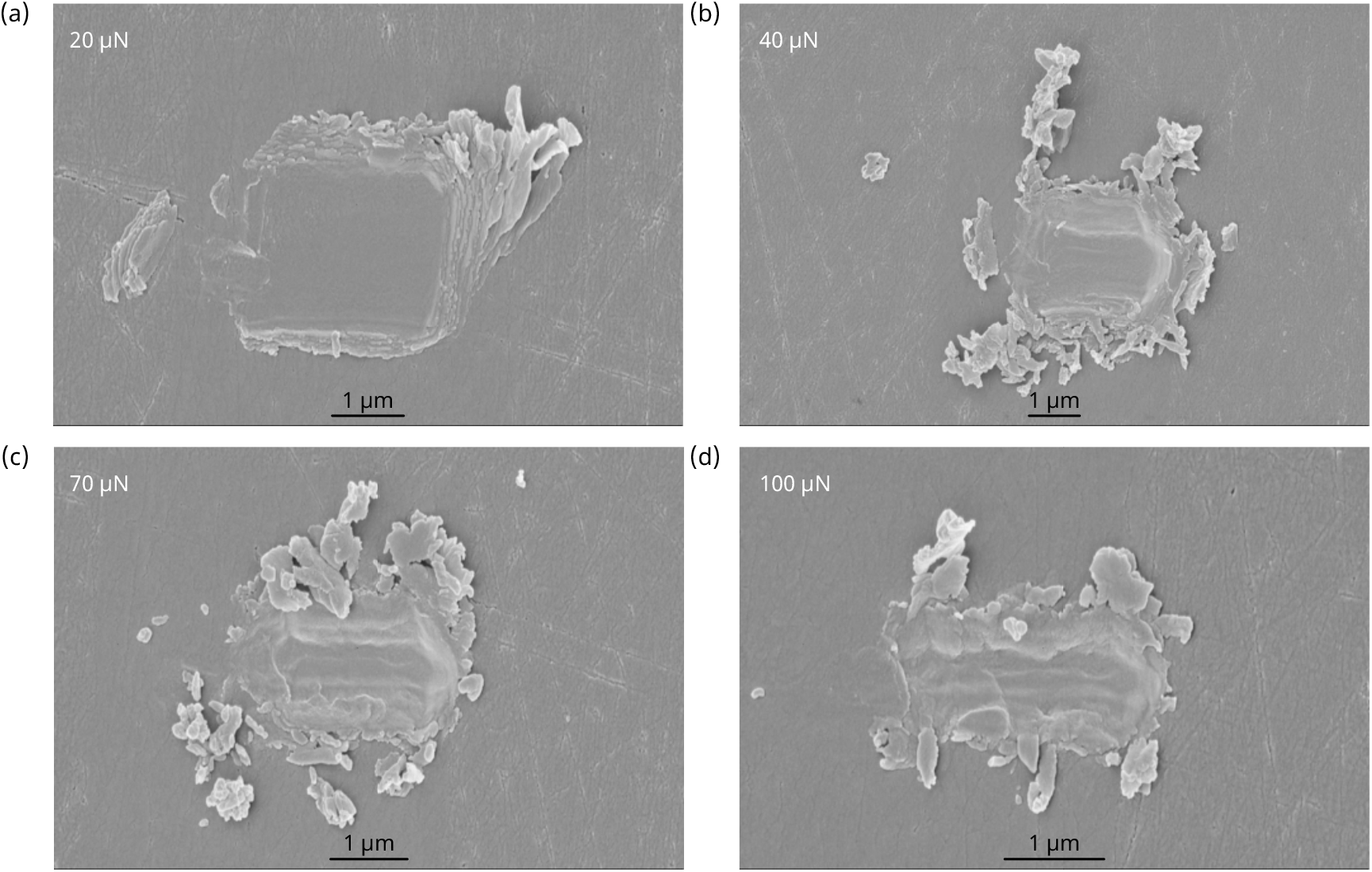
SEM images of nanowear regions. These SEM images were obtained from samples not subjected to post-wear surface topography scanning in order to prevent debris redistribution, for normal wear loads of (a) 20 µN, (b) 40 µN, (c) 70 µN and (d) 100 µN. The relatively smooth surface, absence of obvious cracks, and location and morphology of the wear debris suggest that the wear process is unlikely to be dominated by brittle fracture, abrasion, or microcutting. Rather, it appears more consistent with adhesive wear or delamination.

## 4. Discussion

Mechanical processing of food is an essential task for many insects. Mouth-part wear is the unavoidable consequence of such interactions, and, with time, brings with it potentially severe consequences [19, 28, 25, 18]. But how susceptible are insect mouthparts to wear? To examine how mouthpart material properties contribute to wear resistance, we performed nanowear tests on mandibles from three insect species, at two relative humidities, and under four wear loads. Our results reveal a complex relationship between material properties and wear severity: wear was neither simply proportional to load, nor was it inversely proportional to indentation hardness. Indeed, although hydration consistently reduced epicuticle indentation hardness, it left wear resistance unaffected, or even increased it. In the following sections, we discuss possible origins of these discrepancies between a naive application of classic wear theory and experiments on insect cuticle, and assess the extent to which its nanowear resistance can be linked explicitly to its mechanical properties.

### Indentation hardness is a poor proxy for plastic resistance in insect epicuticle

Wear is a complex phenomenon, as is readily apparent from the engineering literature, which contains upwards of 180 wear models, populated by more than 100 different parameters [128]. Among these models, Archard’s law remains arguably the most popular, owing both to its simplicity and to the clear physical picture from which it is derived [129]. Specifically, a core tenet of Archard-style wear models is the assumption that load is carried by plastically deformed contacts, so that the mean contact pressure—the indentation hardness, *H*_*I*_– is proportional to the material’s yield stress; it is this plastic contact assumption that underpins the predictions that wear severity is inversely proportional to *H*_*I*_, and directly proportional to wear load, *P* [71]. Because our experiments did not bear out these predictions, questioning this assumption appears a reasonable starting point.

A simple qualitative argument can be put forth to illustrate why *H*_*I*_ cannot serve as a universal measure of plasticity for blunt contacts: yielding occurs only above a finite load, below which deformation is fully reversible, and the contact stress field is purely elastic; above that threshold, the mean contact stress is bounded by the yield stress, but some elastic deformation remains; *H*_*I*_ thus generally reflects contributions from both the plastic and elastic properties of the material. Accordingly, the classic Archard scaling— linear in load and inverse in hardness—emerges only asymptotically in the idealised rigid-plastic limit; for real materials, the relationship can be non-linear instead, with the strength of the deviation governed by the elastic contribution to the contact response [130]. To estimate the elastic contribution in the wear contacts studied here, we invoke the lumped-parameter model for the indentation response of an elasto-plastic material proposed by [116]. In this framework, *H*_*I*_ emerges naturally as an elasto-plastic hybrid quantity that depends on the material’s true hardness, *H*_*T*_, its indentation modulus, *E*_*I*_, and the contact geometry:

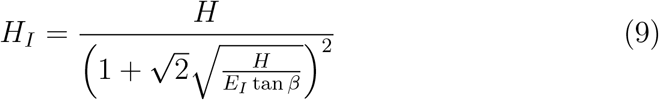

or, equivalently,

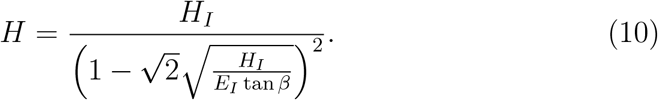

Thus, *H*_*I*_ is a good proxy for *H* only in the limit *H*_*I*_*/E*_*I*_ *→* 0 [116, 131, 62]. For ductile metals, for example, *H*_*I*_*/E*_*I*_ = *O*(10^*−*3^), and the difference between *H*_*I*_ and *H*_*T*_ is modest; for a Berkovich-indentor, tan *β ≈* 0.36, and thus *H*_*T*_ */H*_*I*_ *≈* 1.17. For mandible epicuticle, by contrast, 0.06 *< H*_*I*_*/E*_*I*_ *<* 0.072, so that *H*_*T*_ */H*_*I*_ *≈* 6*−*7. In other words, because its contact response retains a substantial elastic component, *H*_*I*_ is not a faithful measure of plastic resistance, but for the elasto-plastic work required to create a unit volume of residual impression [116, 131]—the relationship between wear severity and hardness becomes more complex.

### An elasto-plastic wear model

To investigate whether an elasto-plastic material model can account for the observed deviations from Archard’s law, we follow previous work and assume that scratching may be viewed as a series of indentations [89, 102]. An Archard-like wear law can then be derived from the elasto-plastic lumped parameter model, by assuming that the residual indentation depth, *h*_*r*_, is proportional to the normalised wear depth, *W*_*d*_ (supplementary S14):

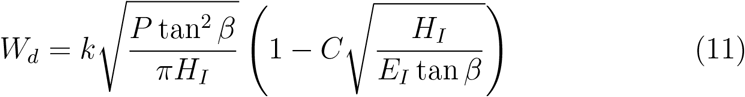

Here, *k* is a dimensionless proportionality constant analogous to Archard’s wear coefficient, whereas the dimensionless constant *C* is related to the yield criterion, and absorbs geometric constants of order unity [116, 132, 131]. For self-similar indenter shapes—those that lack an intrinsic length scale—the face angle *β* is constant, and 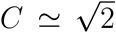. For non-self-similar shapes, however, *β* is load-dependent, and for the cono-spherical tip used in the wear experiments, 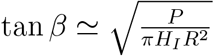, which gives:

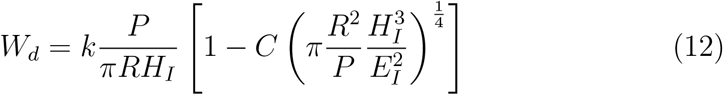

with *R* as the sphere’s radius. Two relevant observations may be extracted from Eq.(12). First, the relation between wear depth and wear load is approximately linear only for loads large compared to the critical load at which yield occurs, 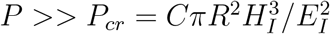[132, 134]; it is non-linear with an exponent larger than unity for all loads below it, and approaches infinity as *P/P*_*cr*_ *→* 1^+^, as revealed by inspection of the derivative:

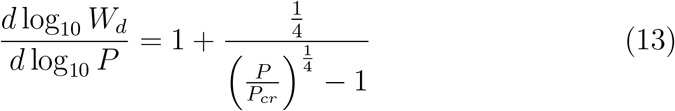

Second, and similarly, the rate of change of *W*_*d*_ with *H*_*I*_ is no longer constant at negative unity:

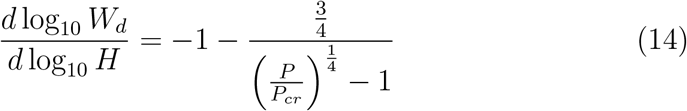

Instead, an inverse proportionality is observed only for *P/P*_*cr*_ *→ ∞*; as *P/P*_*cr*_ *→* 1^+^, the slope decreases, and approaches negative infinity. These predictions are qualitatively consistent with our experimental data (Fig.5 (a-b)), as well as with other empirical studies that reported a relation between wear severity, *H*_*I*_, and *P* that was much stronger than linear [135, 136], attributed to a load-dependent variation of contact strain, and to a breakdown of a direct proportionality between indentation hardness and yield strain [129].

To evaluate the quantitative agreement between the elasto-plastic wear model and our data, we conducted non-linear least-squares regression in MATLAB 2024a (MathWorks, Natick, MA, USA) to fit either Eq.(8) or Eq.(12) to wear data for for both hydration levels, leaving the dimensionless constants *k*_*d*_, *k*, and *C* free (see Fig.5 (e-f) and Fig.7(a-b). See supplementary S12 for the fitted values of the dimensionless constants). Both the Akaike Information Criterion (AIC) and the absolute mean residual error of the elasto-plastic model were consistently smaller, and significantly so at the lowest load (Tab. 4, Fig.7 (c-d)); it thus provided the better fit. Nevertheless, the predictions were not perfect: the elasto-plastic model systematically overpredicted wear at small loads, and underpredicted it at high loads, if less so than the classic Archard law (see Table. 4). As pointed out by [131], it is unlikely that the elastic strain energy is released completely during unloading, as assumed by the lumped parameter model. Instead, some energy likely remains in the impression in form of residual elastic stresses, leading to an underestimation of the residual indentation depth [131]. To account for this effect, [131] proposed an empirical correction. A fit conducted with this up-dated semi-empirical model removes much of the bias at low and high loads, and captures the main trends in the experimental data more accurately (see supplementary S16).

**Figure 7:**
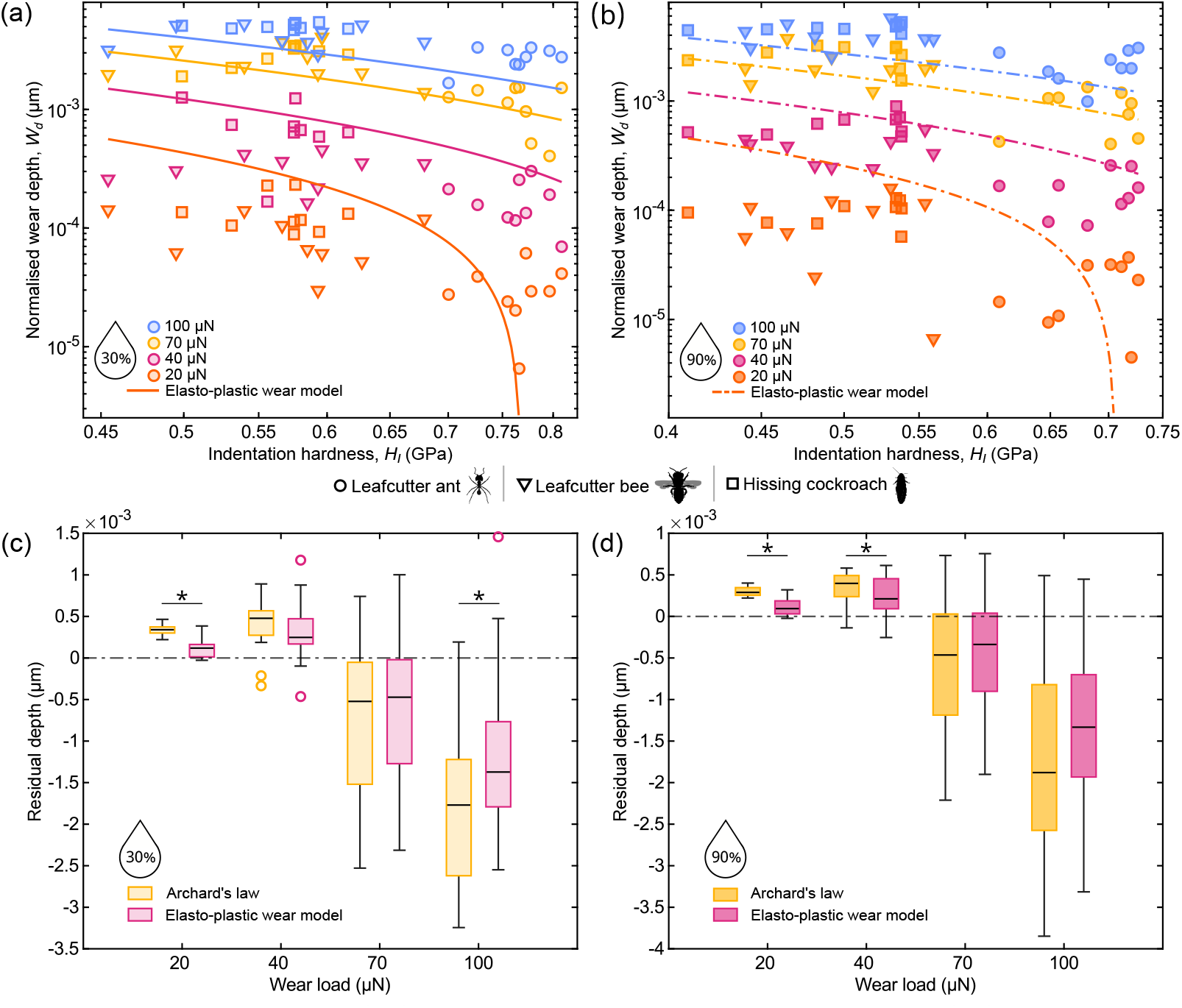
Elasto-plastic wear model fits to the experimental data and residual-error comparison between the two models. Fits of the elastoplastic wear model Eq.(12) to experimental data obtained at all loads by estimating the constants *k* and *C* at (a) 30% and (b) 90% relative humidity (RH). Because the model depends on *E*_*I*_, a continuous function of *H*_*I*_ cannot be constructed; for visual clarity, the species-specific arithmetic mean of the indentation modulus was used instead (for a more complete visualisation, see supplementary S15). Residual errors for the elasto-plastic wear model and Archard’s law Eq.(8) at each normal load are compared for 30% RH (c) and 90% RH (d), which show that the elasto-plastic wear model follows the trends in experimental data significantly better than Archard’s law. In all box plots, the central line indicates the median, the box bounds represent the 25th and the 75th percentile, and whiskers extend to the most extreme data points above and below the median that are within 1.5 times the interquartile range from the box bounds.

**Table 4:** Akaike Information Criterion (AIC) and pairwise comparisons of residual errors from fits of Archard’s law Eq.(8) and the elastoplastic wear model Eq.(12) at 30% and 90% relative humidity (RH).

| load($\mu\text{N}$) | RH | AIC | | $t_{52}$ | p-value |
| --- | --- | --- | --- | --- | --- |
|  |  | Archard’s | elasto-plastic |  |  |
|  |  | law | wear model |  |  |
| 20 | 30% | 451.1 | 414.4 | $W_{52}=1069$ | <0.0001 |
| | 90% | 454.1 | 420.5 | $W_{52}=1061$ | <0.0001 |
| 40 | 30% | 365.5 | 357.6 | $t_{52}=1.49$ | 0.1426 |
| | 90% | 387.7 | 377.4 | $t_{52}=2.11$ | 0.0393 |
| 70 | 30% | 302.3 | 306.5 | $t_{52}=0.89$ | 0.3803 |
| | 90% | 306.9 | 312.8 | $t_{52}=0.58$ | 0.5674 |
| 100 | 30% | 298.8 | 301.4 | $t_{52}=2.51$ | 0.0151 |
| | 90% | 289.9 | 296.6 | $t_{52}=1.53$ | 0.1328 |

### Ashby plots and the search for wear proxies

Wear is difficult to quantify directly, in particular in small biological samples where mass-loss measurements become unreliable. As a result, considerable attention has been devoted to the identification of wear proxies— properties that are more readily measured and that permit at least a qualitative ranking by wear resistance [134, 63, 62, 12, 39, 8]. For yielding, such rankings typically involve indentation hardness and modulus, and different materials are most conveniently compared on Ashby maps that include isoperformance contours [134, 63, 39]. For sharp wear particles, these maps are considered one-dimensional, with contours defined by *H*_*I*_; for blunt wear particles, they become two-dimensional, with indentation modulus as the second axis, and contours defined by 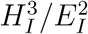[137, 138, 139, 140, 57, 134, 141, 142, 63, 143, 144, 62, 39, 145, 146, 147, 148, 149, 150, Note that *H*_*I*_-contours indicate equal residual indentation depth at a given load, whereas 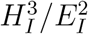 contours indicate equal yield load.].

For elasto-plastic contacts, both maps change fundamentally, and *H*_*I*_ and 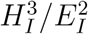 alone no longer define iso-performance contours. The map for sharp contacts becomes two-dimensional, with contours of equal wear severity for a given load given by 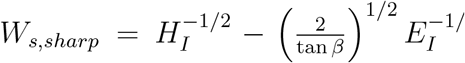 see Fig.8(a) and supplementary S17). For blunt particles with fixed ra-dius *R*, the map becomes three-dimensional, with a third axis formed by the applied wear load, and an iso-performance surface defined by 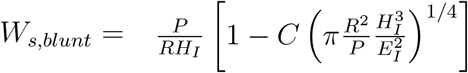 (eq.12 see Fig.8(b) and supplementary S17).For elasto-plastic materials, *H*_*I*_ and 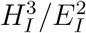 may thus still provide correct qual-itative rankings, but no longer necessarily so. To illustrate this observation with a practical example, we compare hydrated vertebrate enamel [63], hissing cockroach cuticle, and the squid sucker ring [151]. Enamel is about two times harder than hissing cockroach cuticle (1.11 vs 0.50 GPa) and five times harder than the squid sucker rings (1.11 vs 0.20 GPa); the rankings for *H*_*I*_*/E*_*I*_ and 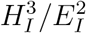, however, are the exact reverse (0.03 vs 0.06 vs 0.09 and 0.013 vs 0.0017 vs 0.0018, respectively). Nevertheless, for sharp Berkovich-like wear contacts, the performance ranking is not correctly predicted by any of the property groups; enamel is predicted to perform the best, hissing cockroach cuticle the worst and squid sucker rings in between (*W*_*s,sharp*_ = 0.535 vs 0.613 vs 0.609 *GPa*^*−*1*/*2^, respectively. Fig.8(a)). In blunt contacts, in turn, all three materials perform similarly at a wear load of 2.4 µN, but enamel performs better than hissing cockroach cuticle at 300 µN (*W*_*s,blunt*_ = 0.662 vs 1438 nm), while squid sucker rings fall behind hissing cockroach cuticle (*W*_*s,blunt*_ = 3566 vs 1438 nm) as shown in Fig.8(b).

**Figure 8:**
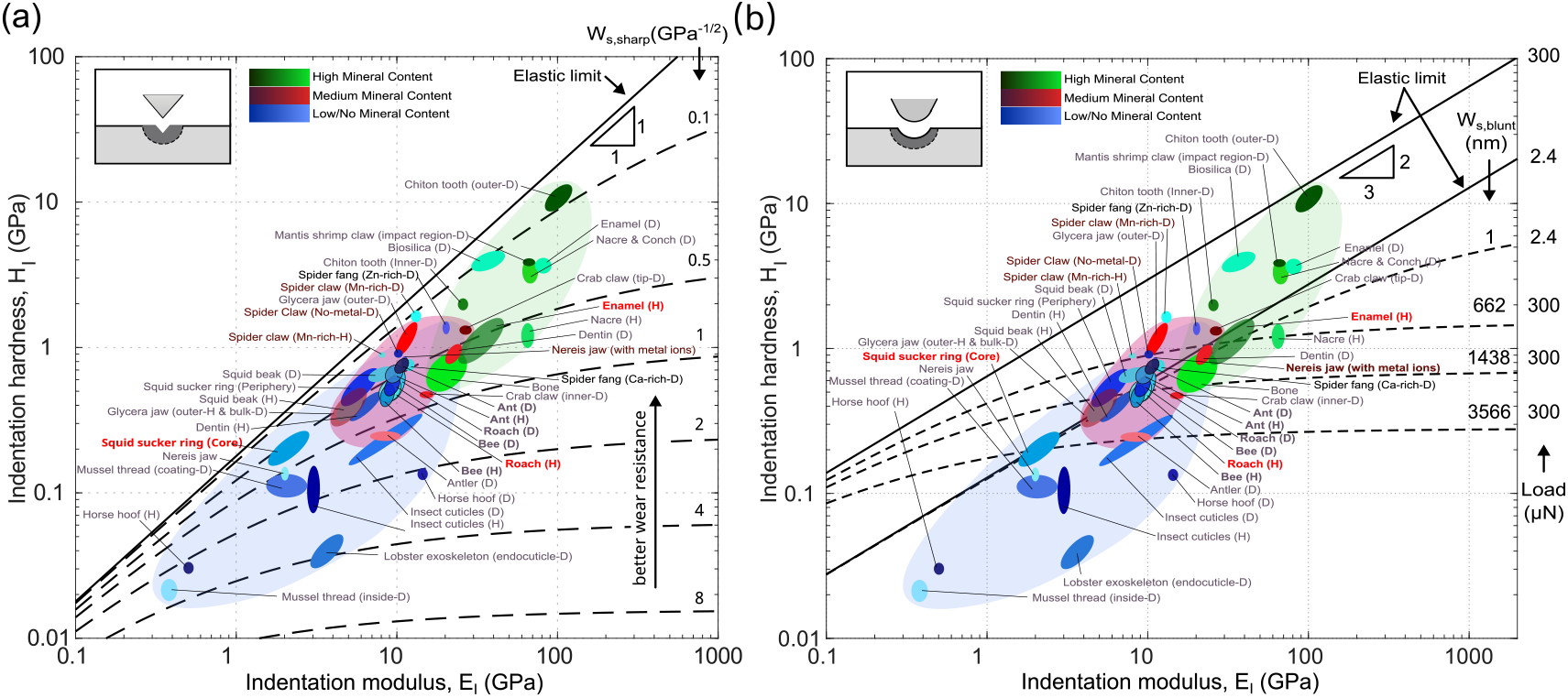
Ashby plots showing iso-performance contours for wear under (a) sharp and (b) blunt contacts; materials that lie on the same line are predicted to have comparable wear performance. In contrast to Archard’s law, which ranks materials by indentation hardness alone, the iso-performance contours for elasto-plastic materials under sharp contact depend on two parameters—indentation hardness and indentation modulus—which together govern the partitioning between elastic and plastic deformation. The solid line indicates a physical limit, corresponding to a material with infinite true hardness and thus to nominally zero wear. For blunt contacts, the situation becomes more complex still, because the ranking of materials can reverse with wear load, because different parameter groups govern resistance to yield initiation, and the partitioning between elastic and plastic deformation once yield has occured. As an illustrative example, vertebrate enamel, hissing cockroach cuticle, and squid sucker ring are predicted to have similar wear performance at low loads, but to diverge at large loads. Note that in both cases, wear coefficients—which can differ by many orders of magnitude across materials but cannot easily be predicted— are assumed to be identical across all materials, and these and similar iso-performance plots should therefore be interpreted with great caution. Data were obtained from Amini & Miserez [63] and Tadayon et al [39].

Swaps in wear rank are likely rare, because both *H*_*I*_ and *E*_*I*_ relate to bond strength and therefore correlate [for biological materials, *H*_*I*_*/E*_*I*_ *≈* 0.05 across many order of magnitude 62]; larger *H*_*I*_ will thus often imply greater predicted wear resistance. However, accounting for elasto-plasticity can still change the magnitude of inferred performance differences. To illustrate this point, we briefly consider the functional significance of transition metals, incorporated in many biological “tools” deployed in mechanically demanding tasks [11, 144, 8, 39, 12], including the mandibles of leaf-cutter ants where they are thought to reduce the wear induced by leaf-cutting [11, 12]. Consistent with this interpretation, leaf-cutter ant mandibles exhibited higher *H*_*I*_, *E*_*I*_, and wear resistance than the mandibles of leaf-cutter bees and hissing cockroaches, which lack transition metals [see also 12]. Notably, however, the differences in wear resistance were larger than expected from *H*_*I*_ alone: ant mandibles were only about 1.3–1.4 times harder, but their wear resistance exceeded that of bee and cockroach mandibles by factors of about 3-4 at low loads, and only approached factors of about 1.5 at the highest wear loads (Fig.4). These differences are consistent with the elasto-plastic wear model, because *E*_*I*_ varied much less among species than *H*_*I*_, leading to a sharper variation in 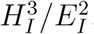. Metal inclusions—as much as other differences in cuticle composition and ultrastructure that may exist between the studied species—may thus be most effective at increasing wear resistance if they affect *H*_*I*_ and *E*_*I*_ differentially.

A more direct assessment of the effect of metal inclusions is enabled by chelation experiments [140, 39]. In *Nereis* jaws and spider claws, removal of metals by chelation reduced *H*_*I*_ by factors of about six and two, respectively [140, 39]. Because the reduction in *E*_*I*_ was smaller, however, the decrease in wear resistance predicted by a sharp elasto-plastic wear model is only a factor of 3 and 1.5, respectively. The contrast is even sharper for blunt contacts, where the classic 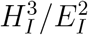 metric suggests a difference of almosta factor twelve for *Nereis* jaws. At sufficiently large loads, however, wear severity is dominated by *H*_*I*_, and the predicted difference drops to a factor of six—roughly half as large.

Of course, the litmus test for any wear proxy is whether it reproduces experimental wear rankings, and here the results are sobering. The largest comparative dataset for biological materials comprises thirteen distinct test regions from seven species [12]—and none of the proxies shows a significant rank correlation with the experimentally determined wear ranks (Holmcorrected Spearman’s *ρ. H*_*I*_, *ρ* = 0.25, 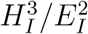, *ρ* = 0.60, *p*=0.125; 1*/W*_*s,sharp*_, *ρ* = 0.32, *p* = 0.88; 1*/W*_*s,blunt*_, *ρ* = 0.27, *p* = 0.88). Perhaps this disagreement stems from a mismatch between the mechanical picture from which wear proxies were derived—an analogy between indentation and scratching—and the experimental assay with which wear was induced—a pin-on-disk geometry [12]. Indeed, in our experiments, where the model resembles the experiment more closely, the relevant blunt-contact proxy performs better: it correctly predicts that ant mandibles outperform those of bees and roaches, and that this advantage diminishes with wear load. Even so, it also predicts similar wear performance for cockroach and bee mandibles, despite the small but consistent experimental advantage of the latter, and—perhaps more concerning—that hydration should reduce wear resistance, when it left it unchanged or even improved it (Fig.4). What explains this discrepancy?

### Cuticle hydration and the problem of wear coefficients

Hydration generally reduces *H*_*I*_ and *E*_*I*_ in biological materials [62], including insect cuticle [39, 11, 7, 90, 63, 84, 38, 91, 94, 95], where water molecules form hydrogen bonds with cuticular proteins [84, 152, 153]. In principle, such changes can decrease or increase the predicted wear resistance, depending on the relative magnitude of the reductions in *H*_*I*_ vs *E*_*I*_ (see above). However, our data and previous work suggest that hydration indeed acts as a “plasticiser” [84], i. e. it reduces the material’s true hardness [Fig.3 62]. What, then, may explain the repeated observation that hydration not only decreases wear volume, but may even improve wear resistance [Fig.4 and 38]?

A likely culprit is the uncertainty in the wear coefficients *k*_*d*_ and *k*—the dimensionless constants in Archard-style wear laws. The problem is simple: wear models typically include a probabilistic component. The wear coefficient, capturing this component, has eluded theoretical prediction and is thus usually included as a proportionality constant that must be determined experimentally. Remarkably, Archard’s wear coefficient may vary by as much as seven orders of magnitude across materials [130], and even within a relatively constrained abrasive-wear regime in metals, it can vary by one to two orders of magnitude [154]. Moderate, let alone small, differences in *H*_*I*_ and *E*_*I*_ are thus unlikely to be reliable indicators of differences in wear resistance, and it should perhaps be expected that wear proxies can perform poorly, even if the underlying models are mechanically sound. The data presented in this manuscript provide a case in point: the elasto-plastic wear model performs equally well within dry and hydrated epicuticles, and the fitting routine even returns statistically indistinguishable yield constants *C* (*Z*=1.27, *p*=0.2010); but the wear coefficient *k* differs significantly by about 30 % (*Z*=3.60, *p*=0.0003), more than enough to offset the difference in wear resistance predicted from material properties. Within dry and hydrated materials, the elasto-plastic model thus performs well enough, perhaps even surprisingly well. But across them it fails, because it cannot predict the wear coefficient *k*.

A change in the wear coefficient can reflect a change in wear mechanism [155, 156, 157, 158, 159]. However, no obvious differences in the optical appearance of the wear regions between dry and hydrated samples were apparent in SEM images. Perhaps the difference in *k* stems from additional energy dissipation arising from viscoelasticity, common in biological materials, and typically more pronounced in hydrated samples [160]; or perhaps it reflects variable importance of delamination wear, [130, 125, 161, 162, 127], in which microimperfections in the material coalesce and grow under repeated low-load cycles, leading to subsurface crack formation, and eventually flake-like wear debris. Clearly, further work is needed to understand the wear mechanisms at play and to link them with material hydration.

### 4.1. Conclusion and outlook

Mandibles perform crucial functions in insects with biting-chewing mouth-parts. While morphological differences among mandibles have been studied extensively [163, 164, 165, 166, 167, 85, 86, 168, 169, 76], our understanding of their mechanical performance—and of its links to morphology and material properties—remains nascent [12, 41, 42, 43, 44, 45, 46, 47, 48, 49, 50, 51, 52, 53]. Limiting mandible wear is likely a crucial aspect of performance [19, 28, 20, 21, 25, 18, 27, 26, 23, 170], but it is difficult to study, because wear is a system property that depends on contact geometry, the properties and shape of the wear bodies, and environmental conditions [54, 55]. In this work, we conducted controlled nanowear experiments designed to loosely resemble the scratching of mandibles by small particles during feeding—one important, though by no means the only; wear mode that mandibles may encounter. We showed that the resistance to such wear can be modelled with fair accuracy using an elasto-plastic material model that treats scratching as analogous to indentation; wear resistance is then load-dependent and depends either on a combination of 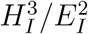 and *H*_*I*_ at low loads, or on *H*_*I*_ at high loads. However,variations in the wear coefficient, not captured by such models, can rival or even exceed the observed variation in material properties, thereby rendering predictions inaccurate or even incorrect. Future work will have to address the wear mechanisms involved in repeated scratching, and design controlled wear experiments that closely resemble other biologically plausible wear scenarios at the nano-, meso-, and macroscale. A cross-disciplinary effort integrating materials science, mechanics, and animal behaviour will be needed to deepen our understanding of how insects may minimise mandible wear—and how plants may best increase it.

## Supporting information

Nanomechanics of Mandible Wear in Insects - SI

## CRediT authorship contribution statement

Dilanka I. Deegala: Conceptualisation, Data curation, Formal analysis, Investigation, Methodology, Validation, Visualisation, Writing - original draft, Writing - review & editing. Jonathan G. Pattrick: Data curation, Methodology, Writing - review & editing. David Labonte: Conceptualisation, Funding acquisition, Investigation, Methodology, Project administration, Resources, Supervision, Visualisation, Writing - original draft, and Writing - review & editing.

## Acknowledgement

This research has received funding from the European Research Council (ERC) under the European Union’s Horizon 2020 Research and Innovation Programme (grant agreement no. 851705, to David Labonte). We acknowledge Maryam Tadayon for sharing data, Liam Crowley for confirming bee identification, and Yael Politi and Marc Masen for insightful discussions that helped improve an earlier version of this manuscript.

