## Supplementary material for "Nanomechanics of Mandible Wear in Insects": Nanomechanics of Mandible Wear in Insects - SI

### S1. Left and right mandible comparison of mature leaf cutter ants

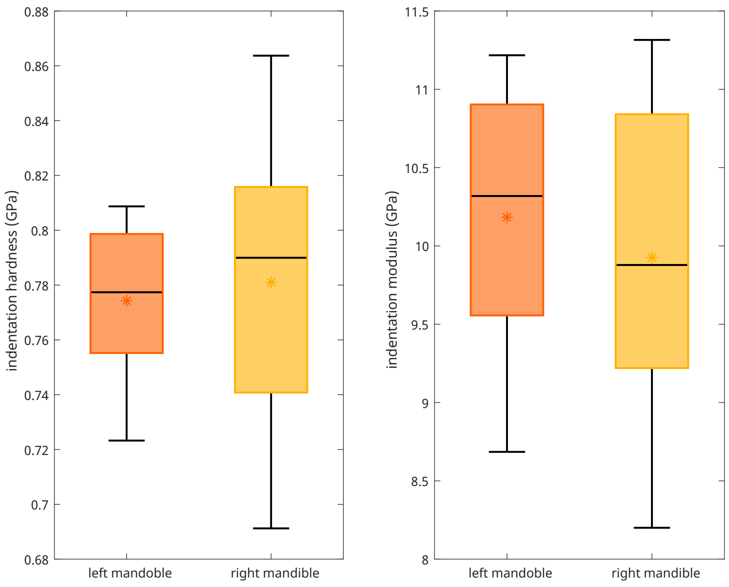


**Figure S1.** Indentation hardness and modulus were not significantly different between left and right mandibles for mature leaf cutter ants. Paired t-tests, Hardness: $t_7$=-0.41 p = 0.6966, Indentation Modulus: $t_7$=1.2678 p = 0.2454.

### S2. Anchor points for the jig

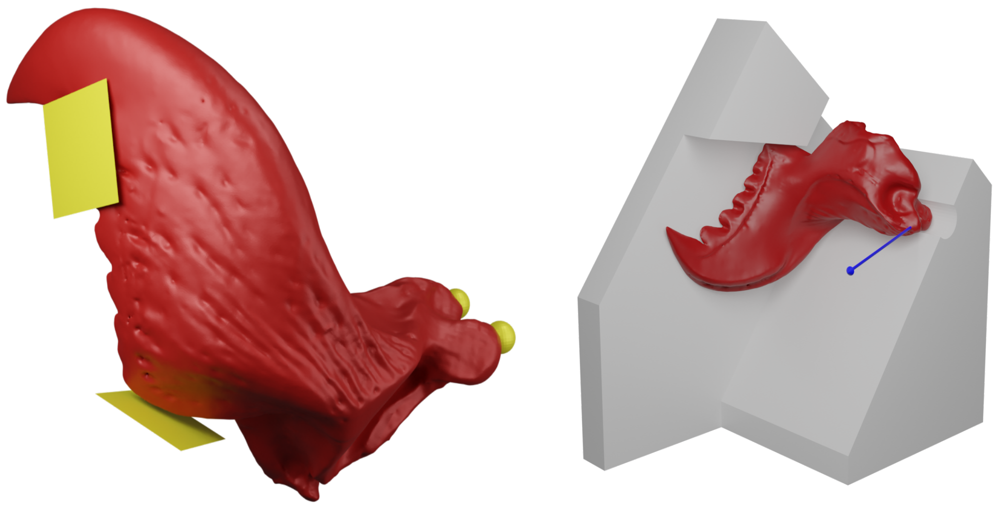

**Figure S2.** left: anchor points are shown in yellow, spheres show the location of the dicondylic joints and the planes indicate the surfaces with less effect from wear, right: a mandible attached the jig, the blue arrow indicates the approximate axis of rotation.

### S3. Property mapping

n=10 leaf-cutter ant mandibles were embedded and polished following the procedure described in the method section. After the first polish, indentations were conducted on the epicuticle of teeth with an exposed crossection with a Berkovitch tip. A 3-step partial unloading load function was used with maximum loads of 1000 µN, 1500µN, and 2000µN. Lower loads were selected to increase the number of indents placed on the epicuticle. After the indentation, samples were repolished with 4000P abrasive paper and polished with 0.3µm alumina powder using a polishing cloth. Each polishing step removed around 30µm of material. As the data collected was not balanced, Scheirer–Ray–Hare test was done on both indentation hardness and indentation modulus and results are shown below.

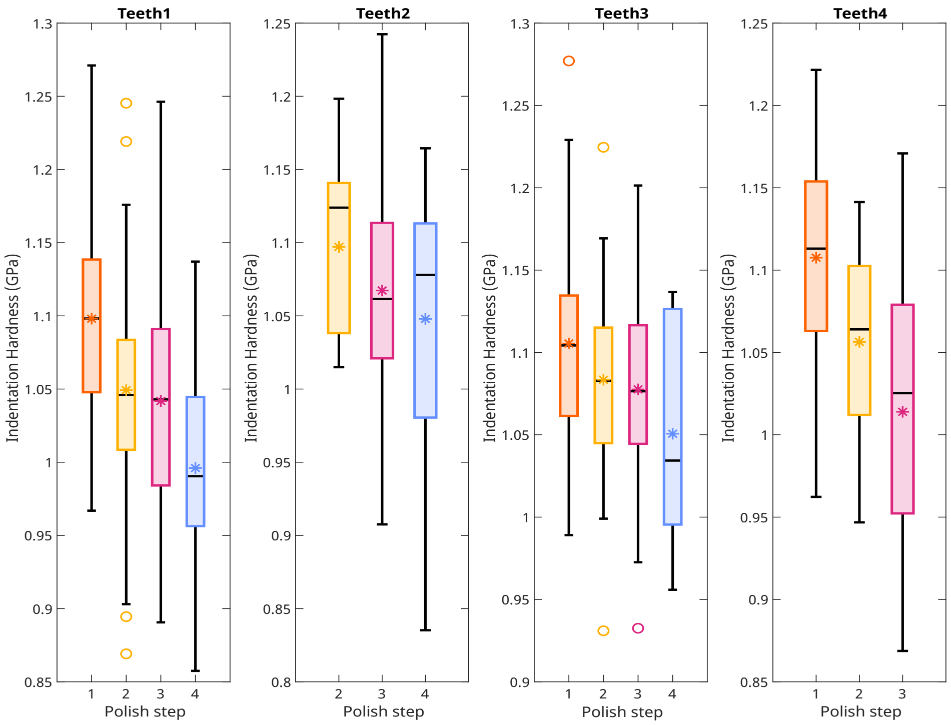


**Figure S3.** Variation of indentation hardness between polish step and teeth. In all box plots, the central line indicates the median, the box bounds represent the 25th and the 75th percentile, and whiskers extend to the most extreme data points above and below the median that are within 1.5 times the interquartile range from the box bounds. Asterisks indicate the means.

**Table S1.** Scheirer–Ray–Hare test results for polish step and teeth on their effect on indentation hardness. Within the test, the response variable was the hardness (continuous variable) and the predictor variables were tooth (categorical variable) and polish step (categorical variable)

| Variable | SS | DF | H | p-value |
|---|---|---|---|---|
| Polish step | 2.03e+06 | 3 | 61.09 | <0.0001 |
| Tooth | 9.66e+05 | 3 | 29.08 | <0.0001 |
| Polish step × Tooth | 2.81e+05 | 7 | 8.46 | 0.2940 |

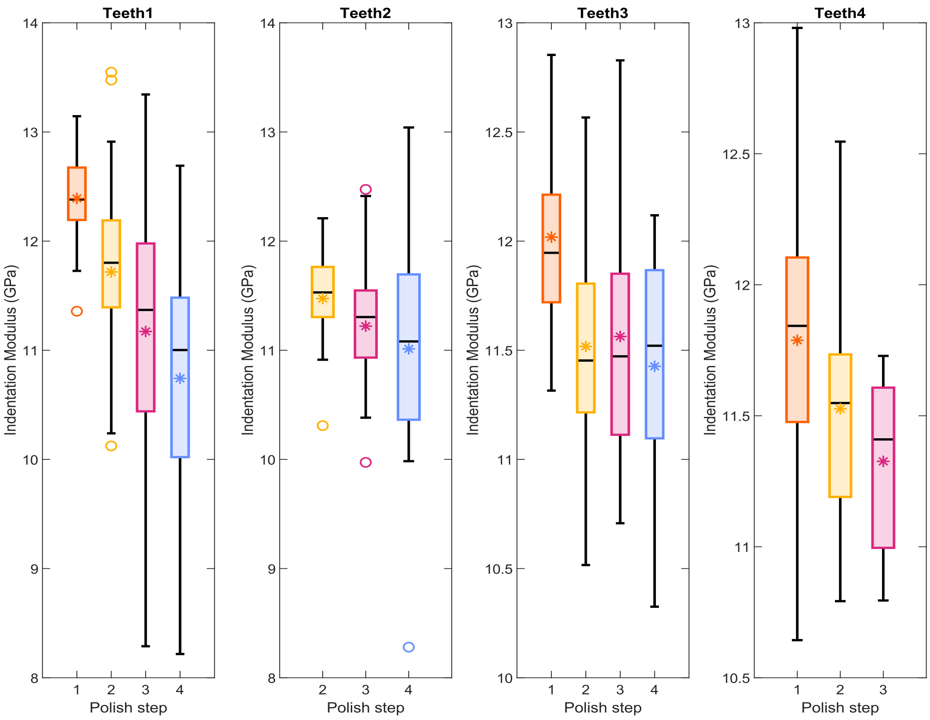


**Figure S4.** Variation of indentation modulus between polish step and teeth. In all box plots, the central line indicates the median, the box bounds represent the 25th and the 75th percentile, and whiskers extend to the most extreme data points above and below the median that are within 1.5 times the interquartile range from the box bounds. Asterisks indicate the means.

**Table S2.** Scheirer–Ray–Hare test results for polish step and teeth on their effect on indentation modulus. Within the test, the response variable was the indentation modulus (continuous variable) and the predictor variables were tooth (categorical variable) and polish step (categorical variable)

| Variable | SS | DF | H | p-value |
|---|---|---|---|---|
| Polish step | 4.12e+06 | 3 | 126.45 | <0.0001 |
| Tooth | 3.17e+05 | 3 | 9.74 | 0.0209 |
| Polish step × Tooth | 6.33e+05 | 7 | 19.41 | 0.0070 |

### S4. Tip area calibration

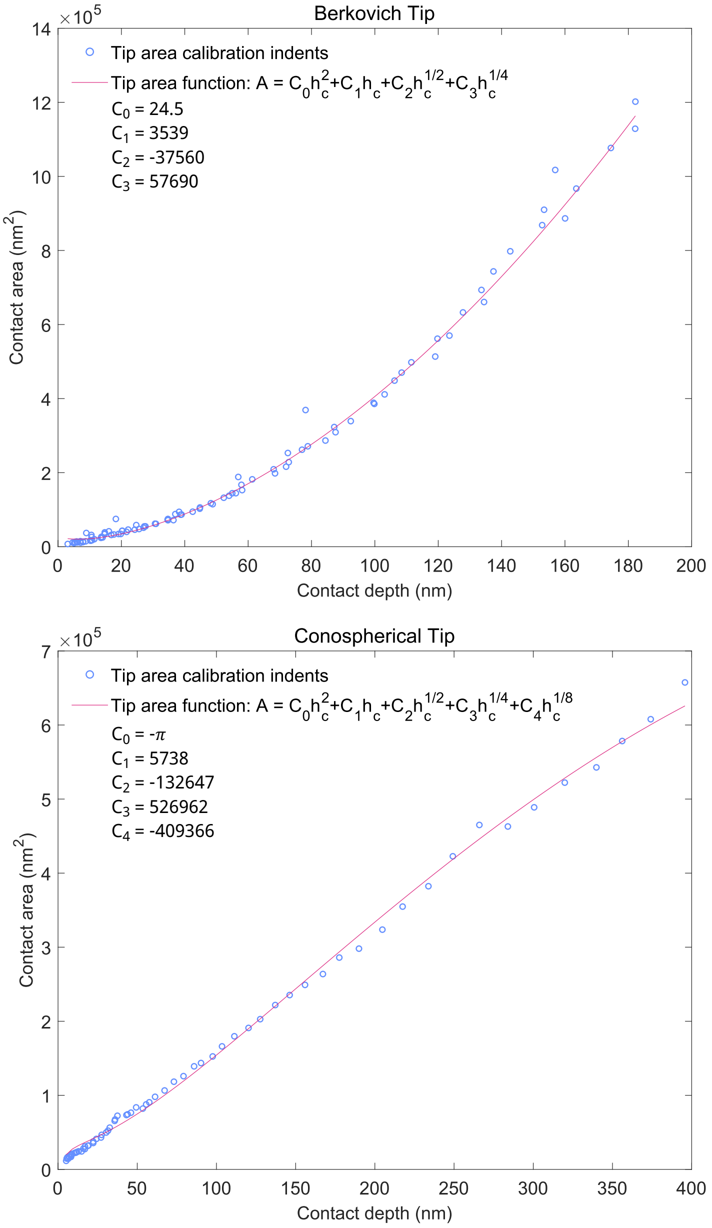


**Figure S5.** Tip area function fitted for calibration data gathered for the Berkovich tip and the Conospherical tip

### S5. Material property differences in the lighter and darker regions

#### *S5.1. Indentation*

Indentations were done in ambient conditions using a cube-corner indenter on two mandibles obtained from 2 individual worker ants in the size range 50-60 mg. The cube corner indenter was used because of the narrow cone angle that allowed indents to be placed closer than a Berkovitch indenter. Standard trapezoidal load function was used with a maximum load of 1500 µN and 5 s hold time. The possible reason for the higher hardness values obtained in these tests compared to those obtained with Berkovich tips is that the geometric contact, which accounts for edge effects in the Oliver-Pharr method, cannot be changed in the Triboscan 10.2.129 software used to extract hardness (default set to 0.75 for Berkovich indenters). Nevertheless, significant differences existed between the groups in both mandibles (Mandible 1: $H(7)=59.26$, $p<0.0001$, Mandible 2: $H(7)=50.15$, $p<0.0001$), and post hoc Wilcoxon tests with Bonferroni corrections showed that group 6 in mandible 1 and group 1 in mandible 2, which were in the lighter regions, had significantly higher hardness than the other regions within their respective mandibles ($p<0.0001$).

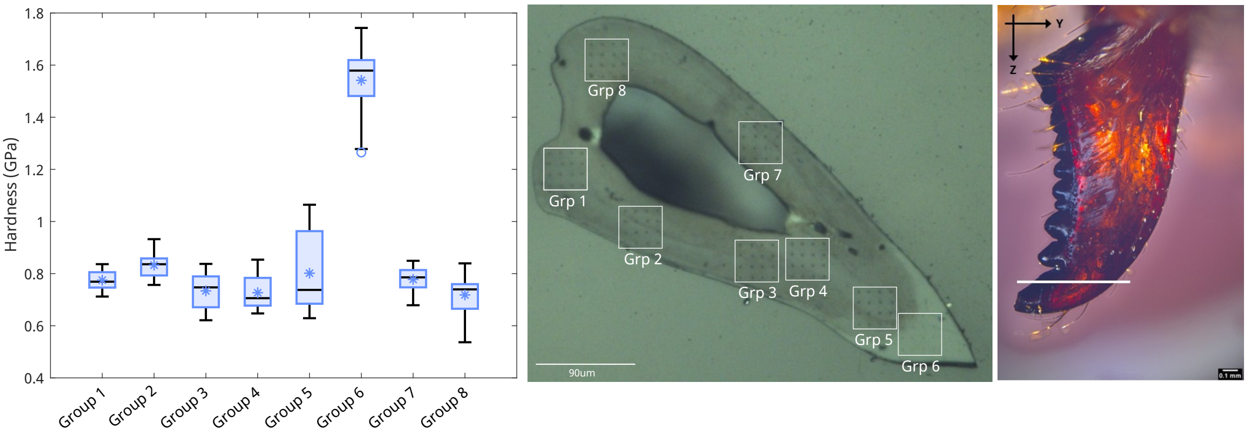


**Figure S6.** Mandible 1. Hardness between the groups was significantly different (Kruskal-Wallis: $H(7)=59.26$, $p<0.0001$). Post-hoc Wilcoxon tests with Benforroni correction revealed that group 6 had significantly higher hardness than the other groups ($p<0.0001$)

#### *S5.2. Scratch tests*

Scratch tests were done on the cuticle with an offset of about 5 µm such that they crossed the optically visible boundary with a cube-corner indenter under a normal force of 100 µm. Two samples were used for this test with cross sections roughly in perpendicular directions, as shown in the figures below. To obtain the frequency constituents, the Lateral force was Fourier transformed with respect to lateral displacement with a custom-written MATLAB code. An upper bound for the frequency range was set at 100Hz to filter out the noise in the signal, and the extracted frequency was 0.8Hz. We expected to identify the presence of layers within the darker regions that cannot be distinguished with optical images, with a characteristic frequency corresponding to the distance between layers. Even though our analysis failed to identify such a distinct frequency, the average of the 10 highest frequency amplitudes in the 0.8-2 Hz range was significantly higher in the darker region compared to the lighter areas (XY-crossection: $t_{58}=5.74$, $p<.0001$, YZ-crossection: $t_{58}=3.27$, $p=.0018$,). This suggested that the lighter region is more homogeneous than the darker region. Mean of the 10 highest frequency amplitudes in the 2-100Hz range were significantly lower than in the range 0.8-2Hz (XY-crossection: $t_{198}=12.26$, $p<.0001$, YZ-crossection: $t_{298}=14.77$, $p<.0001$,).

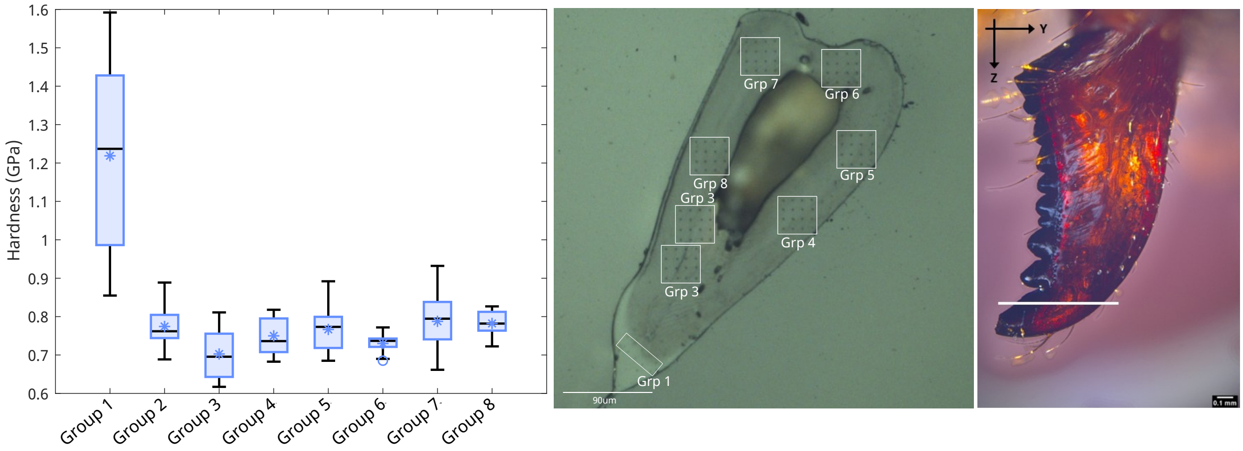


**Figure S7.** Mandible 2. Hardness between the groups was significantly different (Kruskal-Wallis: $H(7)=50.15$, $p<0.0001$). Post-hoc Wilcoxon tests with Benforroni correction revealed that group 1 had significantly higher hardness than the other groups ($p<0.0001$)

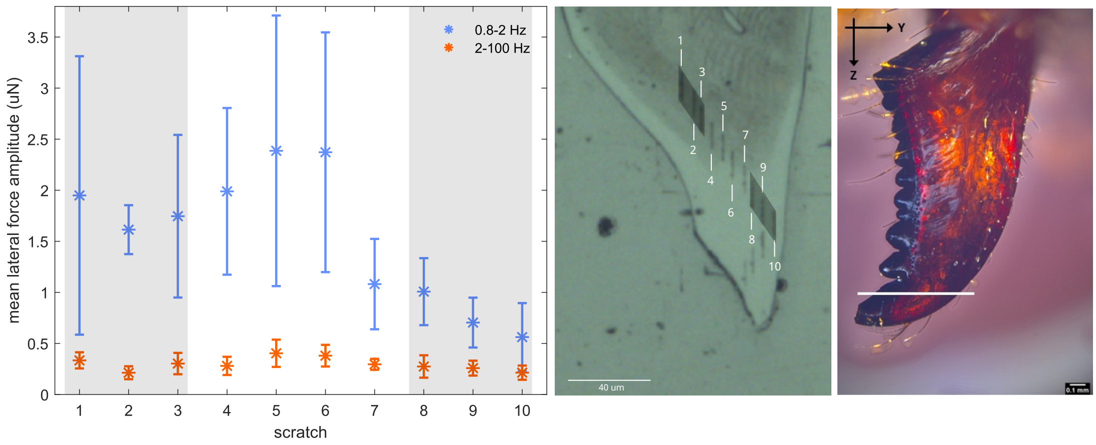


**Figure S8.** XY cross section. Mean of the 10 highest frequency amplitudes in the range 0.8-2 Hz in scratches 1-3 was significantly higher than in scratches 8-10 ($t_{58}=5.74$, $p<.0001$). Mean of the 10 highest frequency amplitudes in the 2-100Hz range were significantly lower than in the range 0.8-2Hz ($t_{198}=12.26$, $p<0.0001$). Error bars indicate the standard deviation.

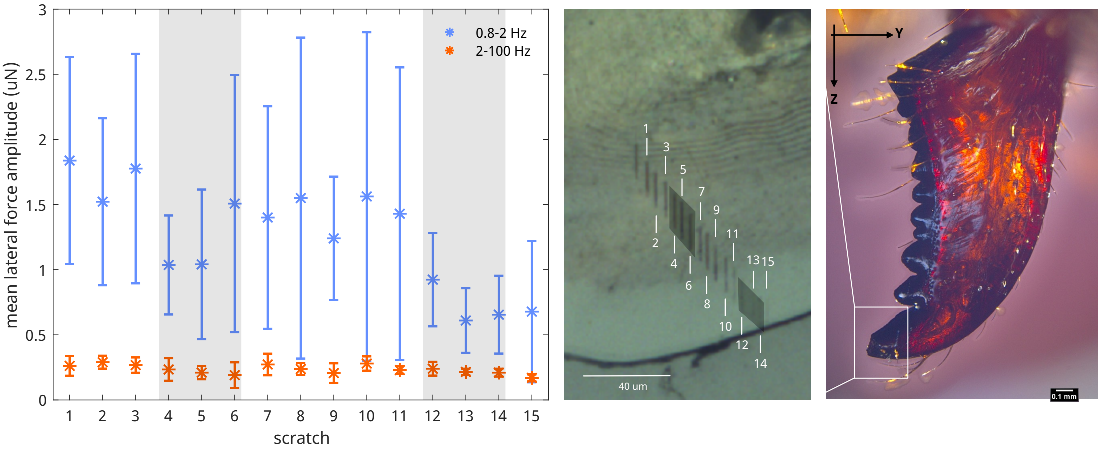


**Figure S9.** ZY cross section. Mean of the 10 highest frequency amplitudes in the range 0.8-2 Hz in scratches 4-6 was significantly higher than in scratches 12-14 ($t_{58}$=3.27, $p$=.0018). Mean of the 10 highest frequency amplitudes in the 2-100Hz range were significantly lower than in the range 0.8-2Hz ($t_{298}$=14.77, $p$<.0001). Error bars indicate the standard deviation.

### S6. Changes in material properties with humidity

Preliminary tests were conducted to measure the variation in properties when humidity is increased from 30%RH to 70%RH and 30%RH to 90%RH. Both indentation hardness and indentation hardness increased significantly only when humidity was increased from 30%RH to 90%RH (H: $t_{14}$=3.80, $p$=0.0020, E: $t_{14}$=4.97, $p$=0.0002)

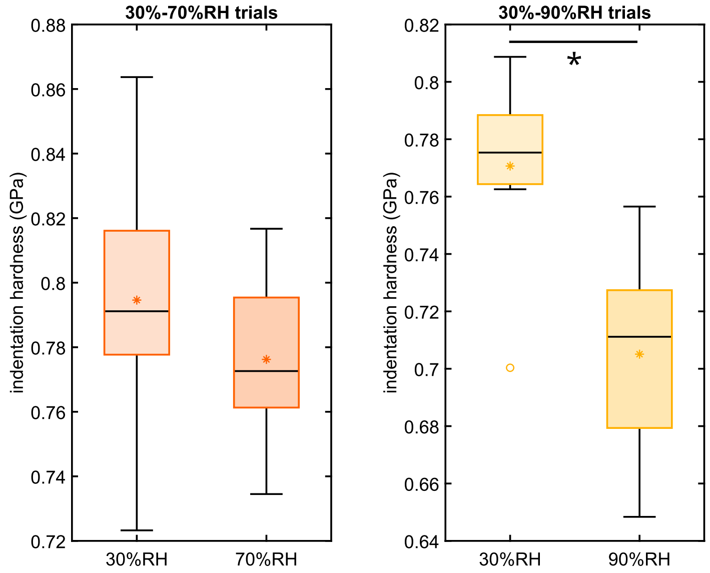


**Figure S10.** Indentation hardness variation with humidity

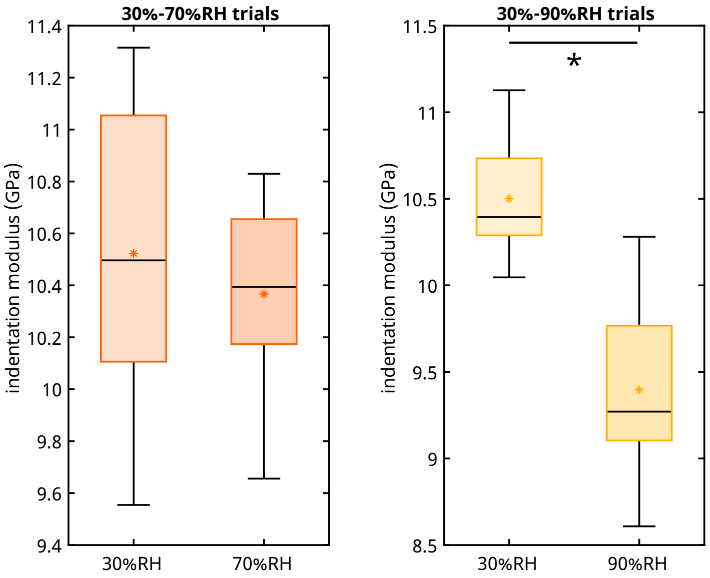


**Figure S11.** Indentation modulus variation with humidity

### S7. Time taken for submerged samples to reach steady state

Mature ant samples were prepared with the same method as mentioned and were submerged using a similar setup to [1]. Epicuticle properties were measured using nanoindentation using a Berkovitch fluid cell tip for 6 hours in 30-minute intervals. This interval was selected because of the extended time period required to find the sample surface and reduce drift measurements. Measurements were also taken after 24hrs and 48hrs of the same samples and shown in following figures. It can be seen that the sample required around 5 hours to reach the same levels as the measurements taken after 24hrs.

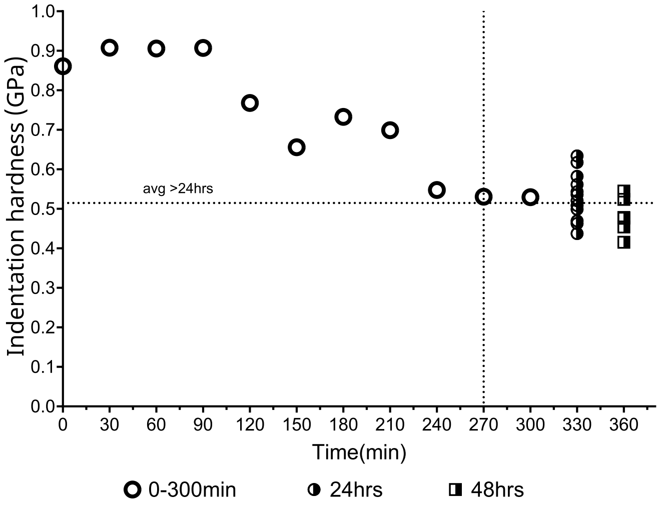


**(a)** Variation in indentation hardness with time

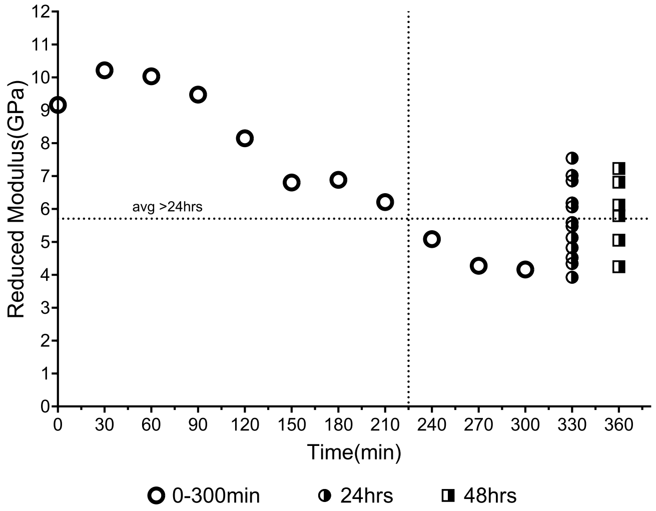


**(b)** Variation in indentation modulus with time

**Figure S12.** Indentation hardness and modulus variation with submerged time

### S8. Variation of Indentation hardness and modulus with time at 90%RH

Indentation hardness and modulus measured during two test runs are plotted against time, with the first conducted on four mandibles and the second on six mandibles. Linear regression did not indicate a significant negative slope in any of the tests, indicating that the steady state was reached during the tests.

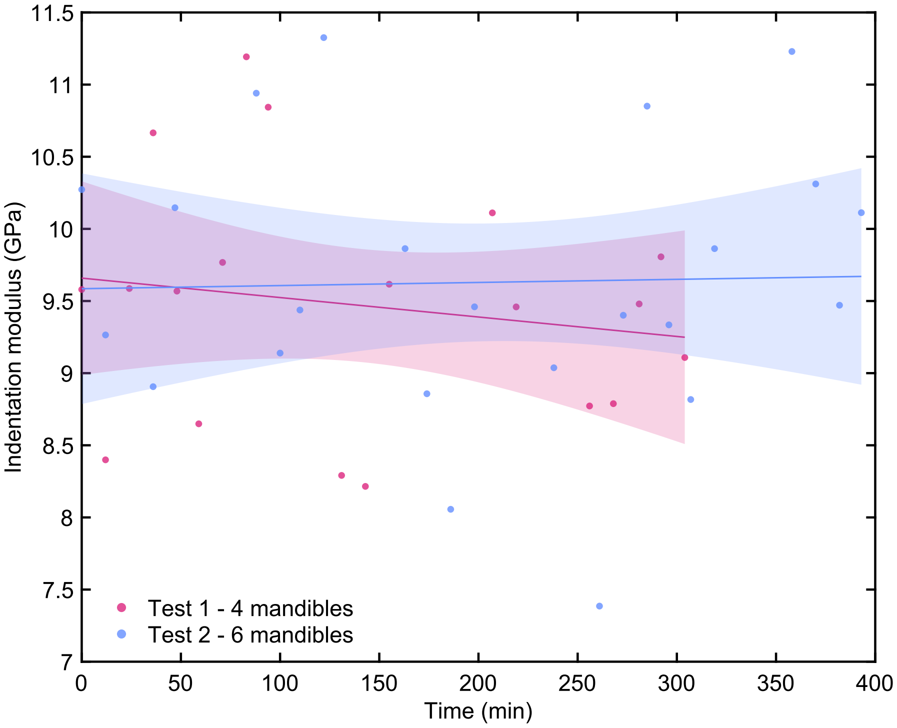


**Figure S13.** Variation of Indentation modulus with time at 90%RH

**Table S3.** Gradient of linear regression between indentation modulus and time at 90%RH. 95% confidence intervals are provided in brackets

| | gradient | t | DF | p-value |
|---|---|---|---|---|
| Test 1 - 4 mandibles | -1.3E-03 [-5.2E-03, 2.5E-03] | -0.7372 | 17 | 0.4710 |
| Test 2 - 6 mandibles | 2.16E-04 [-3.1E-03, 3.6E-03] | 0.1343 | 21 | 0.8945 |

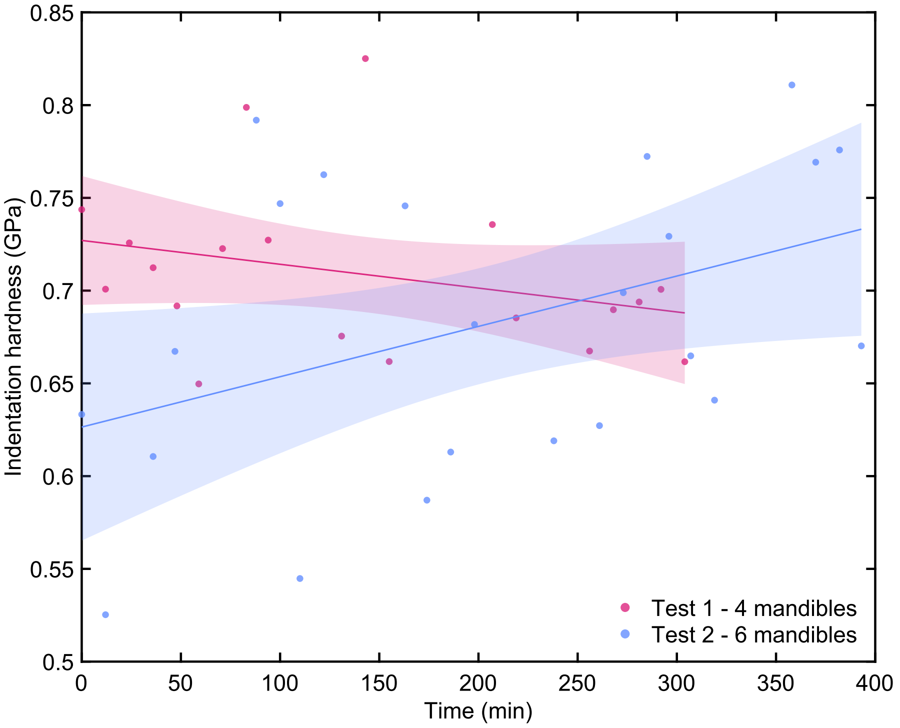


**Figure S14.** Variation of Indentation hardness with time at 90%RH

**Table S4.** Gradient of linear regression between indentation hardness and time at 90%RH. 95% confidence intervals are provided in brackets

| | gradient | t | DF | p-value |
|---|---|---|---|---|
| Test 1 - 4 mandibles | -1.29E-04 [-3.28E-04, 7.14E-05] | -1.3559 | 17 | 0.1929 |
| Test 2 - 6 mandibles | 2.71E-04 [1.48E-05, 5.28E-04] | 2.1994 | 21 | 0.0392 |

#### S9. Sphere-to-cone transition depth

As the manufacturer's datasheet provided, it was assumed that the indenter takes a spherical shape below the transition depth and a conical shape above. The optimisation problem was set up such the shape of the indentor was described by two geometries below and above the transition depth. Indentation modulus was calculated below the transition depth, with the contact area calculated from a spherical contact, and above that depth, from a truncated cone. The indentation modulus obtained in this manner was compared to the manufacturer-provided standard sample reduced modulus. The square mean of the residual was minimised using an Evolutionary algorithm implemented in Microsoft Excel by varying the transition depth, truncated height, radius, and cone angle. This resulted in a radius of 300 nm and transition depth of 102 nm.

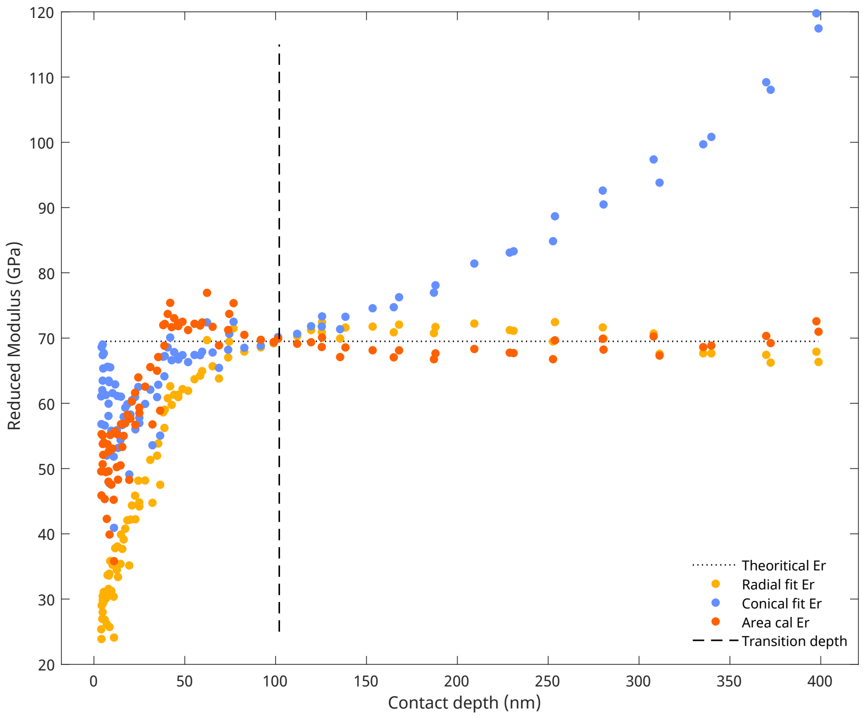


**Figure S15.** Sphere to cone transition depth

### S10. wear depth increased approximately linearly with passes

Wear tests were conducted with a normal load of 100 µm with a scan area of 6 µm×6 µm for 5 passes on 4 different leaf cutter ant mandibles. Multiple linear regression resulted in $R^2$ of 0.98, which indicated a good agreement with linear approximation.

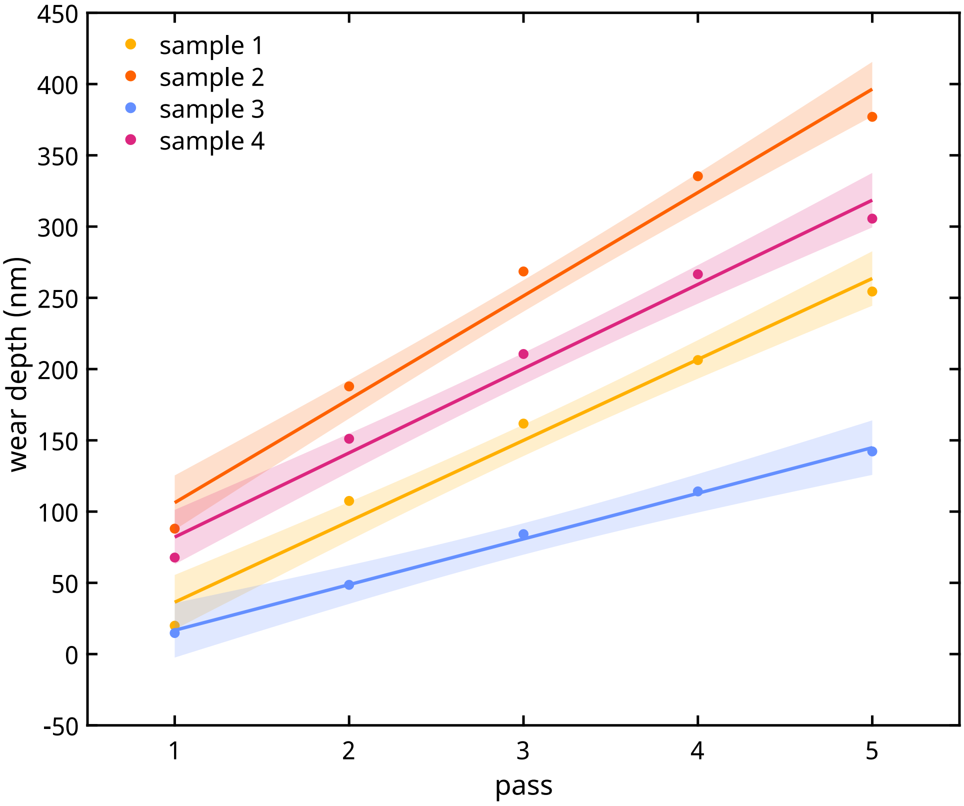


**Figure S16.** Multiple regression resulted in a $R^2$ of 0.98 which indicated a good agreement with linear assumption

### S11. Archard's model wear depth derivation

Here, $V$ is the volume removed during the wear test, $F_N$ the applied normal load, $H$ the hardness of the material, $S(L)$ slide distance per pass as a function of side length $L$, surface area of the worn volume $A(L)$, $h$ the total wear depth, $W_h$ the wear depth per pass, and $K$ is a dimensionless constant representing a probability.

$$V = K\frac{F_N}{H}S(L) \times \#passes$$
$$h \times A(L) = K\frac{F_N}{H}S(L) \times \#passes$$
$$h = K\frac{F_N}{H} \times \frac{S(L)}{A(L)} \times \#passes$$
$$h_{perpass} = W_h = K\frac{S(L)}{A(L)}\frac{F_N}{H}$$

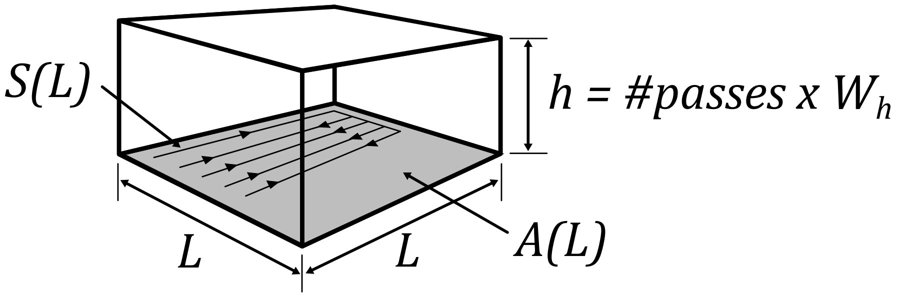


**Figure S17.** Schematic for Archard's model wear depth derivation

### S12. Results of ANOVA

**Table S5.** Results of repeated measures ANOVA conducted to investigate the effect of mandible group and hydration on indentation modulus

| | SS | MS | DF1 | DF2 | F value | p-value |
|---|---|---|---|---|---|---|
| Mandible group | 9.53E+00 | 4.77E+00 | 2 | 24.00 | 14.59 | 0.0001 |
| Humidity | 9.73E+00 | 9.73E+00 | 1 | 24.00 | 74.56 | <.0001 |
| Mandible group × Humidity | 3.33E-01 | 1.67E-01 | 2 | 24.00 | 1.28 | 0.2973 |

**Table S6.** Pairwise comparison of indentation modulus between mandible groups

| Comparison | DF | Q | adj-p-value |
|---|---|---|---|
| Mature ant vs Bee | 24 | 6.11 | 0.0007 |
| Mature ant vs Cockroach | 24 | 7.03 | 0.0001 |
| Bee vs Cockroach | 24 | 0.92 | 0.7941 |

**Table S7.** Results of repeated measures ANOVA conducted to investigate the effect of mandible group and hydration on indentation hardness

| | SS | MS | DF1 | DF2 | F value | p-value |
|---|---|---|---|---|---|---|
| Mandible group | 4.32E-01 | 2.16E-01 | 2 | 24.00 | 76.09 | <.0001 |
| Humidity | 6.81E-02 | 6.81E-02 | 1 | 24.00 | 49.40 | <.0001 |
| Mandible group × Humidity | 3.77E-04 | 1.89E-04 | 2 | 24.00 | 0.14 | 0.8727 |

**Table S8.** Pairwise comparison of indentation hardness between mandible groups

| Comparison | DF | Q | adj-p-value |
|---|---|---|---|
| Mature ant vs Bee | 24 | 15.13 | <.0001 |
| Mature ant vs Cockroach | 24 | 15.09 | <.0001 |
| Bee vs Cockroach | 24 | 0.03 | 0.9997 |

**Table S9.** Results of repeated measures ANOVA conducted to investigate the effect of mandible group and hydration on true hardness

| | SS | MS | DF1 | DF2 | F value | p-value |
|---|---|---|---|---|---|---|
| Mandible group | 79.11 | 39.55 | 2 | 24.00 | 91.47 | <.0001 |
| Humidity | 4.26 | 4.26 | 1 | 24.00 | 23.00 | .0001 |
| Mandible group × Humidity | 0.01 | 0.01 | 2 | 24.00 | 0.03 | 0.9677 |

**Table S10.** Pairwise comparison of indentation hardness between mandible groups

| Comparison | DF | Q | adj-p-value |
|---|---|---|---|
| Mature ant vs Bee | 24 | 16.70 | <.0001 |
| Mature ant vs Cockroach | 24 | 16.43 | <.0001 |
| Bee vs Cockroach | 24 | 0.27 | 0.9801 |

**Table S11.** Results of repeated measures ANOVA conducted to investigate the effect of mandible group, normal load and hydration on wear depth

| | SS | MS | DF1 | DF2 | F value | p-value |
|---|---|---|---|---|---|---|
| Mandible group | 5.34E-05 | 2.67E-05 | 2 | 24 | 98.90 | <.0001 |
| Normal load | 4.75E-04 | 1.58E-04 | 3 | 72 | 600.60 | <.0001 |
| Humidity | 2.00E-06 | 2.00E-06 | 1 | 24 | 9.11 | 0.0059 |
| Mandible group × Normal load | 3.37E-05 | 5.62E-06 | 6 | 72 | 21.33 | <.0001 |
| Mandible group × Humidity | 4.65E-08 | 2.33E-08 | 2 | 24 | 0.11 | 0.8999 |
| Normal load × Humidity | 1.53E-06 | 5.08E-07 | 3 | 72 | 3.58 | 0.0179 |
| Mandible group × Normal load × Humidity | 5.80E-07 | 9.66E-08 | 6 | 72 | 0.68 | 0.6658 |

**Table S12.** Pairwise comparison of wear depth between species at each load

| | load ($\mu$N) | DF | Q | adj-p-value | wear depth ratio |
|---|---|---|---|---|---|
| Ant×Bee | 20 | 24 | 7.51 | <.0001 | 3.2 |
| | 40 | 24 | 4.63 | 0.0086 | 2.1 |
| | 70 | 24 | 8.26 | <.0001 | 2.4 |
| | 100 | 24 | 9.00 | <.0001 | 1.6 |
| Ant×Cockroach | 20 | 24 | 11.73 | <.0001 | 4.5 |
| | 40 | 24 | 13.42 | <.0001 | 4.1 |
| | 70 | 24 | 10.96 | <.0001 | 2.8 |
| | 100 | 24 | 13.65 | <.0001 | 2.0 |
| Bee×Cockroach | 20 | 24 | 4.23 | 0.0169 | 1.4 |
| | 40 | 24 | 8.78 | <.0001 | 2.0 |
| | 70 | 24 | 2.70 | 0.1580 | 1.2 |
| | 100 | 24 | 4.65 | 0.0084 | 1.2 |

**Table S13.** Pairwise comparison of wear depth between hydration levels at each load

| load ($\mu$N) | DF | Q | adj-p-value |
|---|---|---|---|
| 20 | 24 | 2.32 | 0.1135 |
| 40 | 24 | 0.99 | 0.4892 |
| 70 | 24 | 3.57 | 0.0185 |
| 100 | 24 | 3.20 | 0.0331 |

**Table S14.** Pairwise comparison of wear depth between mandible groups

| Comparison | DF | Q | adj-p-value |
|---|---|---|---|
| Ant vs Bee | 24 | 12.92 | <.0001 |
| Ant vs Cockroach | 24 | 19.56 | <.0001 |
| Bee vs Cockroach | 24 | 6.64 | 0.0003 |

**Table S15.** Numerical values of the wear model constants obtained obtained from fitting the models to the experimental data. 95% confidence intervals are provided in parentheses.

| Hydration | $k_d$ | $k$ | $C$ |
|---|---|---|---|
| 30% | 0.0782 (0.0664, 0.0901) | 0.0117 (0.0103, 0.0132) | 2.6191 (2.5891, 2.6491) |
| 90% | 0.0605 (0.0510, 0.0701) | 0.0083 (0.0072, 0.0095) | 2.6667 (2.6002, 2.7331) |

### S13. $h_r/h_{max}$ comparison

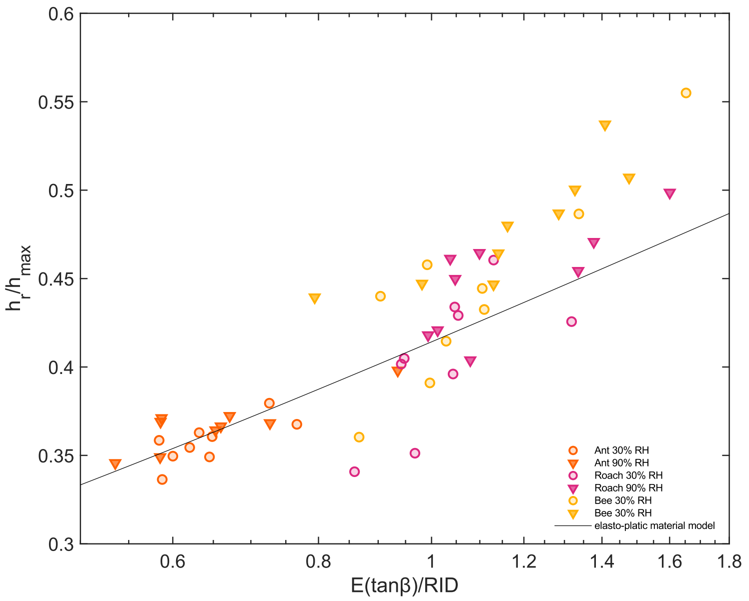


**Figure S18.** $h_r/h_{max}$ comparison between the elasto-plastic material model and nanoindentation data

### S14. Elasto-plastic lumped parameter model

From Sakai1999 Eq.(14a)

$$\frac{h_r}{h_{max}} = \frac{1}{1+\sqrt{2}\sqrt{\frac{H}{E\tan\beta}}} \tag{1}$$

From Sakai1999 Eq.(16)

$$H_I = \frac{H}{\left[1+\sqrt{2}\sqrt{\frac{H}{E\tan\beta}}\right]^2} \tag{2}$$

Substituting $x = \sqrt{\frac{H}{E\tan\beta}}$ into eq.(2)

$$\frac{H_I}{E\tan\beta} = x^2\left(1+\sqrt{2}x\right)^{-2} \tag{3}$$

solving for $x$

$$x = \sqrt{\frac{H_I}{E\tan\beta}} = \frac{1}{\sqrt{\frac{E\tan\beta}{H_I}}-\sqrt{2}} \tag{4}$$

from eq.(1) & eq.(4)

$$\frac{h_r}{h_{max}} = 1-\sqrt{2}\sqrt{\frac{H}{E\tan\beta}} \tag{5}$$

as $h_{max} = \frac{\tan\beta}{\sqrt{\pi}}\sqrt{\frac{P}{H}}$

$$h_r = \frac{\tan\beta}{\sqrt{\pi}}\sqrt{\frac{P}{H}}\left(1-\sqrt{2}\sqrt{\frac{H}{E\tan\beta}}\right) \tag{6}$$

### S15. Elasto-plastic and classic Archard's model fit to experimental data with a variable *E*

Elasto-plastic wear model fit without using an average reduced modulus. (a) model fits for 30%RH, (b) model fits for 90%RH

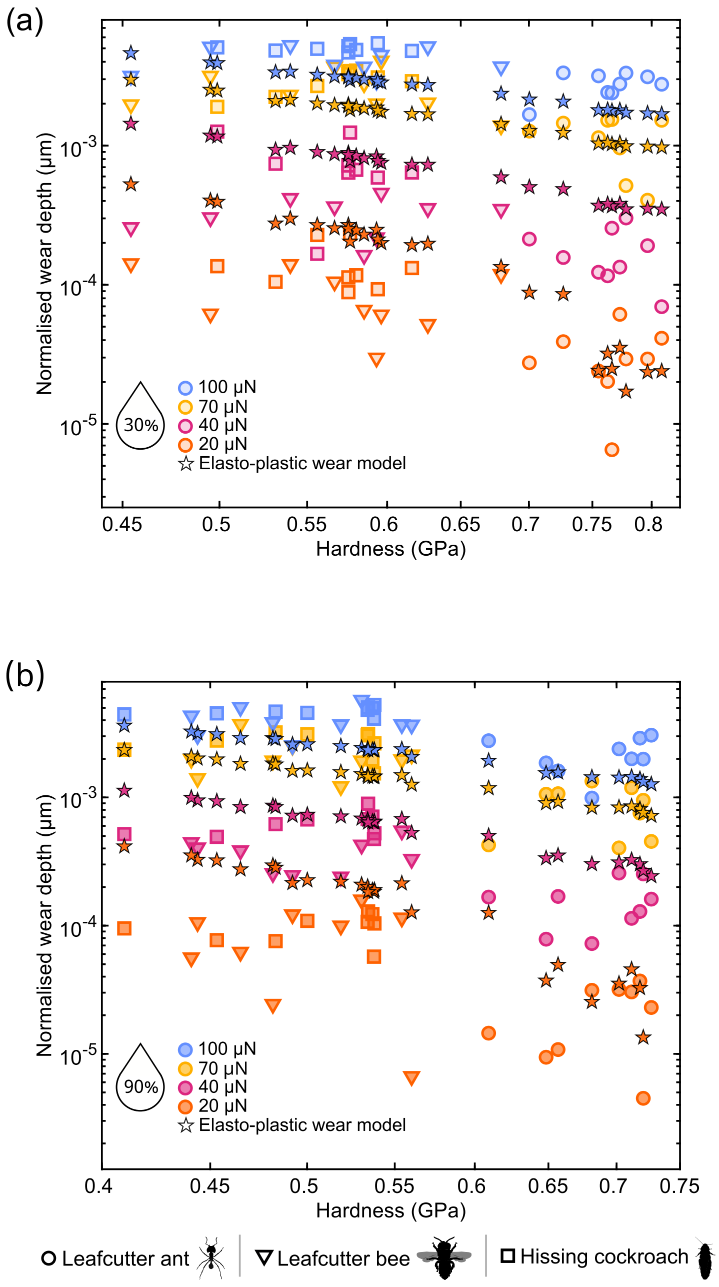


**Figure S19.** Elasto-plastic wear model fits with variable $E$

### S16. Elasto-plastic semi-empirical wear model fits

Sakai et al. 2004 [2] introduced the following elasto-plastic semi-empirical model that accounts for the effect of residual stresses in the deformed material and therefore fits better than the model presented earlier [3], Eq.(1).

$$\frac{h_r}{h_{max}} = \frac{1}{1 + 2\frac{H}{E\tan\beta}} \tag{7}$$

However, the $\frac{h_r}{h_{max}}$ predicted by the elasto-plastic semi-empirical model differed significantly from the values obtained from nanoindentation force displacement data ($t_{103.49}$=8.35, $p$<0.0001) as shown below.

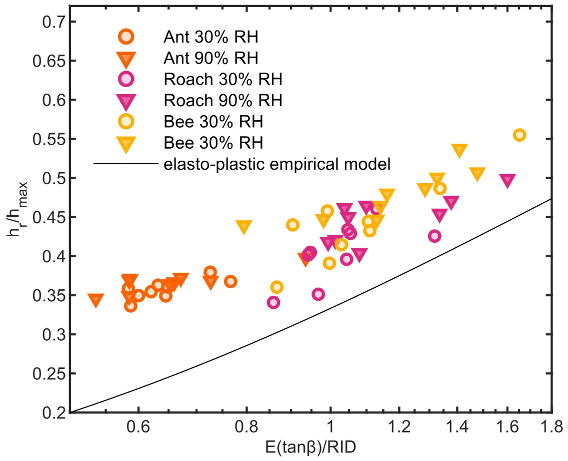


**Figure S20.** Elasto-plastic semi-empirical model fit

Nevertheless we derived a wear model using Eq.(7) following similar steps as in S14, which provided the following where $k_e$ and $C_e$ are dimensionless constants of order unity,

$$W_d = k_e \frac{P}{\pi R H_I} \left[ \frac{1}{1 + C_e \sqrt{\pi \frac{R^2}{P} \frac{H_I^3}{E_I^2}} \left( \sqrt{C_e \sqrt{\pi \frac{R^2}{P} \frac{H_I^3}{E_I^2}}} - 1 \right)^{-2}} \right] \tag{8}$$

Similar steps followed to fit and evaluate the elasto-plastic wear models were also followed here, and resulted in the following.

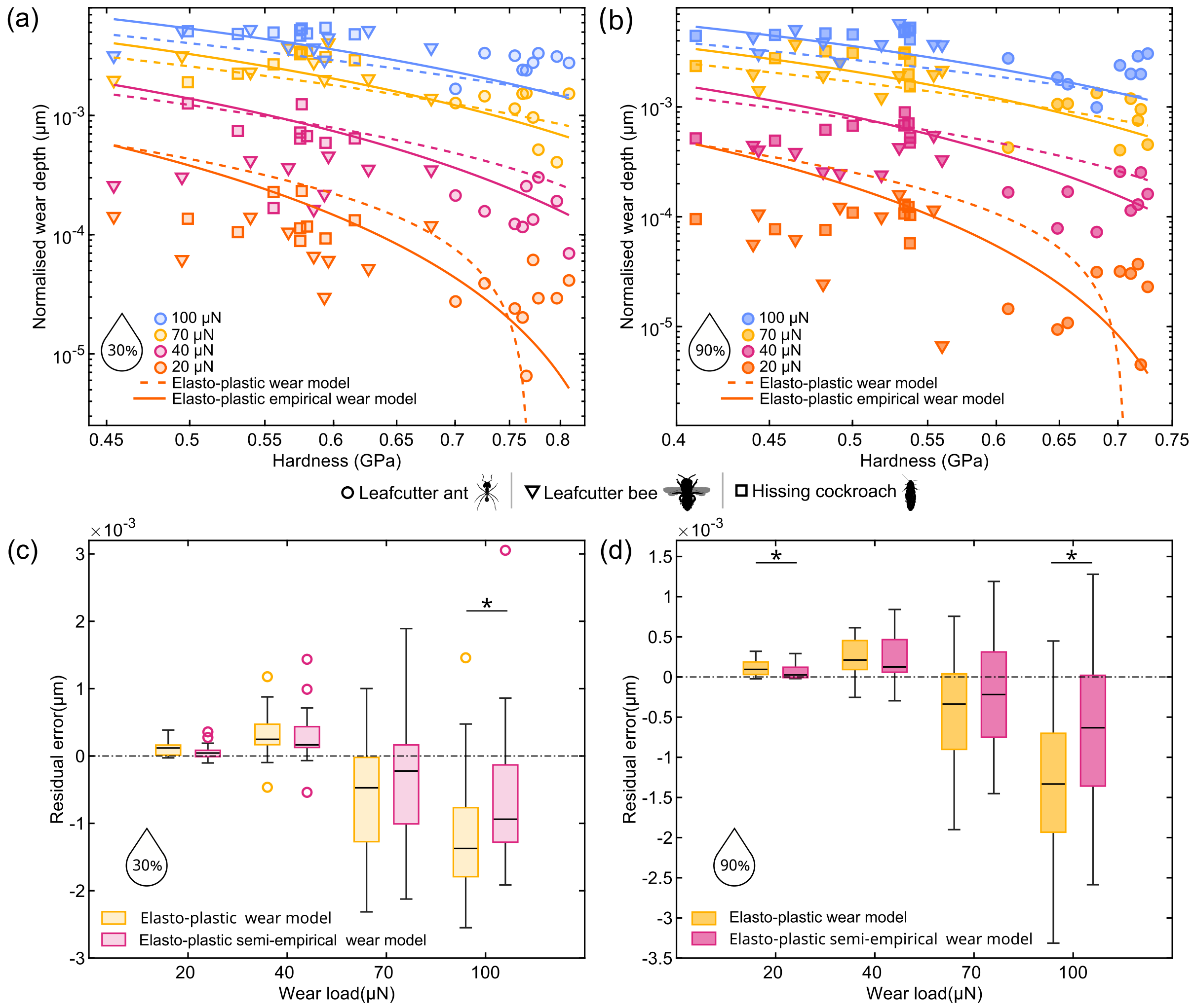


**Figure S21.** Elasto-plastic semi-empirical model fit

The results indicate that although Eq.(7) models do not fit well to indentation data, the elasto-plastic semi-empirical wear model Eq.(8) captures the main trends in the data better than the elasto-plastic wear model.

**Table S16.** Akaike Information Criterion (AIC) and results of pairwise comparison of the residual errors obtained from the elasto-plastic wear model and elasto-plastic empirical wear model Eq.(8) fit to experimental data for 30%RH and 90%RH (relative humidity)

| load($\mu$N) | RH | AIC | | *statistic* | p-value |
|---|---|---|---|---|---|
| | | elasto-plastic empirical wear model | elasto-plastic wear model | | |
| 20 | 30% | 414.4 | 420.5 | $t_{52}$=1.81 | 0.0755 |
| | 90% | 420.5 | 425.3 | $W_{52}$=625 | 0.0430 |
| 40 | 30% | 357.6 | 348.7 | $t_{52}$=0.47 | 0.6416 |
| | 90% | 377.4 | 362.9 | $t_{52}$=0.22 | 0.8303 |
| 70 | 30% | 306.5 | 304.1 | $t_{52}$=0.88 | 0.3839 |
| | 90% | 312.8 | 312.7 | $t_{52}$=1.22 | 0.2277 |
| 100 | 30% | 301.4 | 293.5 | $t_{52}$=2.18 | 0.0332 |
| | 90% | 296.6 | 296 | $t_{52}$=2.3 | 0.0254 |

### S17. Additional Ashby plots for sharp and blunt contacts

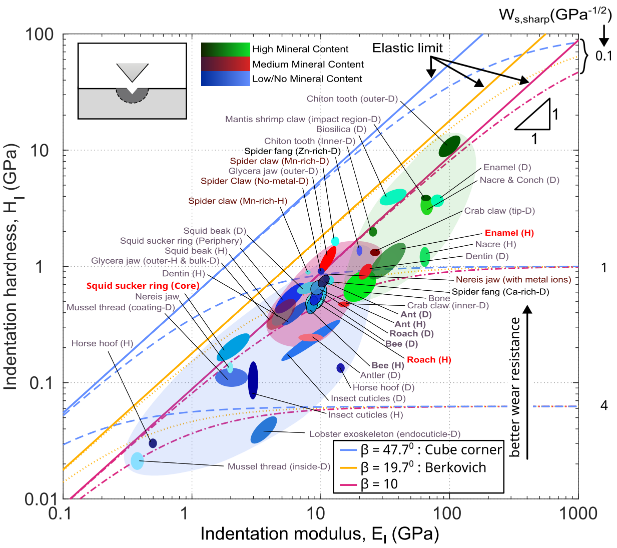


**Figure S22.** Ashby plot with elasto-plastic wear proxy for three indentors that vary in the equivalent cone angle

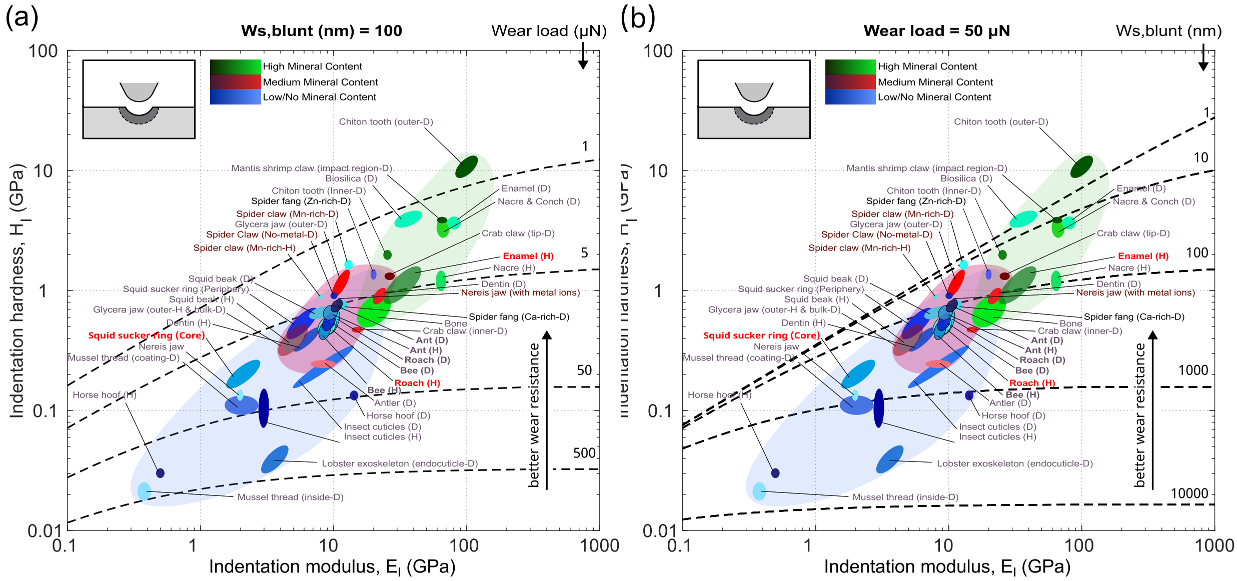


**Figure S23.** Ashby plots with blunt contact elasto-plastic wear proxy. (a) Variation in iso-performance lines under variable load conditions (b) Variation in iso-performance lines under constant load
